# Inherited resilience to clonal hematopoiesis by modifying stem cell RNA regulation

**DOI:** 10.1101/2025.03.24.645017

**Authors:** Gaurav Agarwal, Mateusz Antoszewski, Xueqin Xie, Yash Pershad, Uma P. Arora, Chi-Lam Poon, Peng Lyu, Andrew J. Lee, Chun-Jie Guo, Tianyi Ye, Laila Barakat Norford, Anna-Lena Neehus, Lucrezia della Volpe, Lara Wahlster, Diyanath Ranasinghe, Tzu-Chieh Ho, Trevor S. Barlowe, Arthur Chow, Alexandra Schurer, James Taggart, Benjamin H. Durham, Omar Abdel-Wahab, Kathy L. McGraw, James M. Allan, Ruslan Soldatov, Alexander G. Bick, Michael G. Kharas, Vijay G. Sankaran

**Author notes:** Correspondence: V.G.S., M.G.K.

## Abstract

Somatic mutations that increase hematopoietic stem cell (HSC) fitness drive their expansion in clonal hematopoiesis (CH) and predispose to blood cancers. Although CH frequently occurs with aging, it rarely progresses to overt malignancy. Population variation in the growth rate and potential of mutant clones suggests the presence of genetic factors protecting against CH, but these remain largely undefined. Here, we identify a non-coding regulatory variant, rs17834140-T, that significantly protects against CH and myeloid malignancies by downregulating HSC-selective expression and function of the RNA-binding protein MSI2. By modeling variant effects and mapping MSI2 binding targets, we uncover an RNA network that maintains human HSCs and influences CH risk. Importantly, rs17834140-T is associated with slower CH expansion rates in humans, and stem cell MSI2 levels modify ASXL1-mutant HSC clonal dominance in experimental models. These findings leverage natural resilience to highlight a key role for post-transcriptional regulation in human HSCs, and offer genetic evidence supporting inhibition of MSI2 or its downstream targets as rational strategies for blood cancer prevention.

## Introduction

It is increasingly recognized that ‘healthy’ aging tissues often harbor a substantial burden of cancer driver mutations, underscoring the widespread nature of somatic mosaicism^1^. In the hematopoietic system, myeloid malignancy (MyM)-associated mutations are commonly detected in the peripheral blood of otherwise healthy individuals, a phenomenon termed clonal hematopoiesis (CH). The current paradigm is that a subset of randomly acquired mutations in hematopoietic stem and progenitor cells (HSPCs) confer a fitness advantage^2^, with pervasive positive selection in aging individuals causing increasing CH driven by relatively few HSC clones^3,4^. Those with detectable CH of indeterminate potential (CHIP), defined based on the presence of MyM-associated single nucleotide variants above ∼2% in the peripheral blood, have an elevated lifetime risk of developing MyM^5^ and a range of chronic disorders^6–8^, with a causal role demonstrated in some cases such as cardiovascular disease^6,9^. However, while there has been significant progress in describing the natural history and consequences of CH, the mechanisms driving dominance of stem cell clones remain poorly understood.

Recently, mechanistic investigation based on genome-wide association studies (GWAS)^10–14^ have identified genetic regulators of stem cell clonality. For example, functional follow-up of inherited MyM risk variants have revealed increased HSC self-renewal and transcriptional elongation driving MyM predisposition^11,14–17^. In contrast, the germline mechanisms protecting HSCs from CH/MyM remain poorly understood. Increasing depth of sequencing has demonstrated that CH driver mutations are observed ubiquitously in aging adults^18^, yet most never progress to a large clone size or to an overt MyM. Moreover, lineage tracing suggests that driver mutations may be acquired in early life, decades prior to disease, suggesting there may be mechanisms promoting disease latency^19,20^. Indeed, longitudinal profiling^3,21–23^ and inference of clonal expansion rates^17^ show that the growth rate and potential of mutant HSC clones varies significantly. While external factors such as chronic inflammation^24^ and chemical exposures^25^ could contribute to this variable expansion, these observations suggest that germline genetic variation might also promote resilience to CH/MyM in some individuals.

Here, we leverage human genetic variation to identify an inherited mechanism protecting HSCs from CHIP, through downregulation of the RNA-binding protein MSI2. By faithfully modeling variant effects in primary human hematopoietic cells and conducting mechanistic studies, we illuminate an MSI2-regulated RNA network that is important for human HSC maintenance and which is attenuated through genetic variation to resist stem cell expansion and progression to MyM.

## Results

### Identification of a CHIP and MyM resilience haplotype at the 17q22 locus

Given population variability in the expansion of mutant CH clones^3,21–23^, we hypothesized that there may be inherited mechanisms protecting individuals from developing a large clone size and MyM. To explore this, we conducted a GWAS meta-analysis for CHIP. We replicated associations at 24 loci reported previously [in UK Biobank (UKB) and Geisinger Health Study (GHS)]^11^ in the All of Us (AoU) cohort and conducted a pooled meta-analysis of 43,619 CHIP-carriers and 598,761 controls (**Fig 1A, Supp Table 1**). This analysis identified rs80093687 at the 17q22 locus – a common single nucleotide polymorphism (SNP) across populations (**Supp Fig 1A-B**) – as among the most protective variant associated with CHIP-resilience [additive OR=0.84, 95% CI=0.82-0.87, p=9.6×10^-22^] (**Fig 1B**), with homozygotes carrying an ∼30% reduced risk for CHIP (**Fig 1C, top**). Notably, an independent GWAS of CHIP-carriers^26^ that involved 3 additional cohorts had replicated the protective 17q22 association (**Supp Fig 1C**), with their lead SNP in high linkage disequilibrium (LD) with rs80093687 [R^2^=0.87] (**Supp Fig 1D**), suggesting a shared CHIP-protective haplotype at this locus, where the underlying mechanisms have remained unexplored.

**Figure 1:**
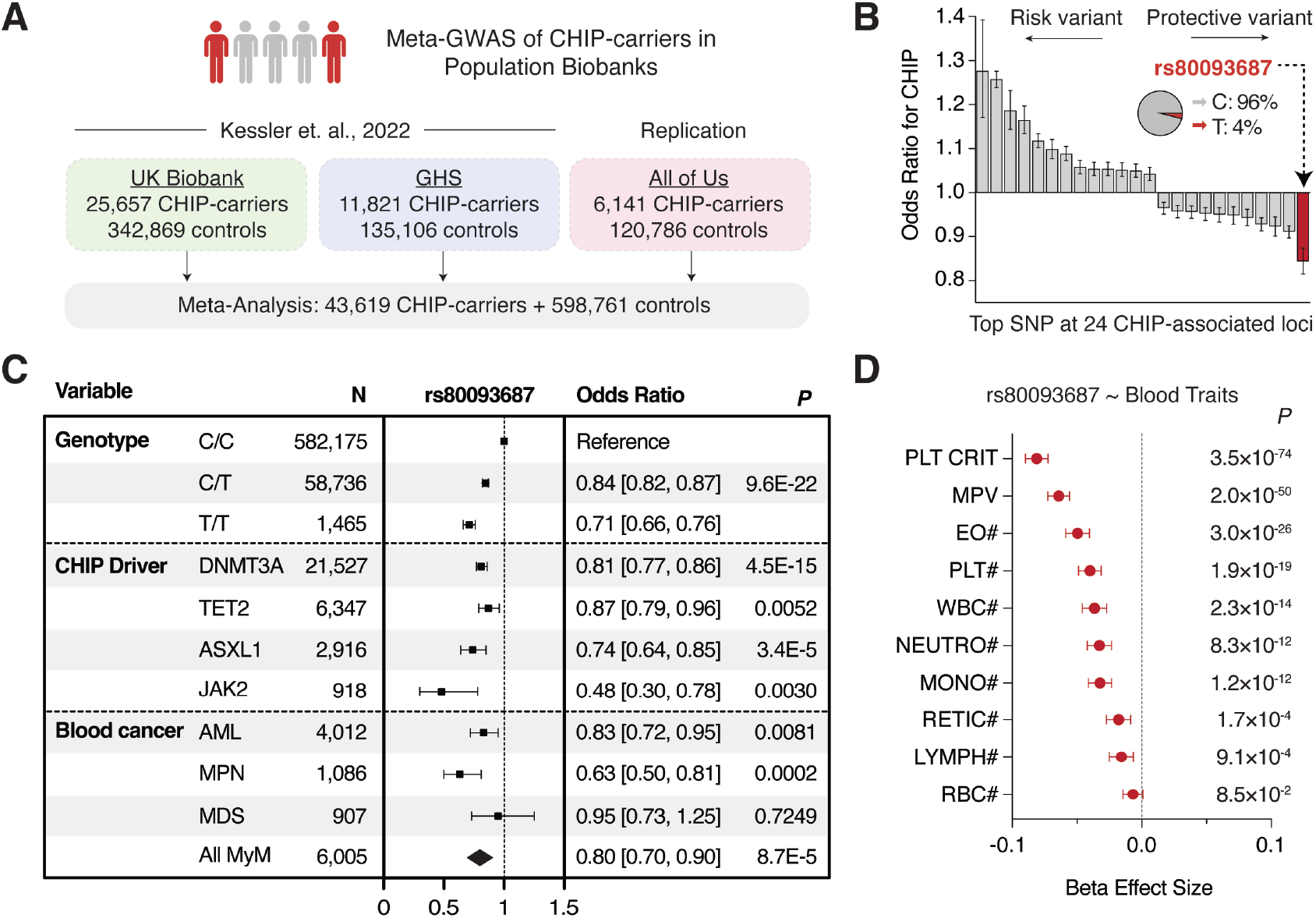
Inherited resilience to CHIP and myeloid malignancy at the 17q22 locus. **(A)** Approach for meta-analysis of clonal hematopoiesis of indeterminate potential (CHIP) across UK Biobank (UKB), Geisinger Health Study (GHS) and All of Us (AoU) cohorts. **(B)** Waterfall plot showing meta-analysis odds ratio effect sizes (95% CI) for the most significant variant at 24 CHIP-associated loci. **(C)** Protective effect of rs80093687 for CHIP by genotype (top), by driver mutation (middle), or for myeloid malignancies (MyMs) (bottom). AML = acute myeloid leukemia; MPN = myeloproliferative neoplasm; MDS = myelodysplastic syndrome. **(D)** Associations between rs80093687 and blood cell traits. PLT CRIT = plateletcrit; MPV = mean platelet volume; EO = eosinophil; PLT = platelet; WBC = white blood cell; NEUTRO = neutrophil, MONO = monocyte; RETIC = reticulocyte; LYMPH = lymphocyte; RBC = red blood cell; # = count.

In a meta-analysis, rs80093687 was associated with resilience to several canonical CHIP driver mutations, including *DNMT3A*-CHIP [OR=0.81], *TET2*-CHIP [OR=0.87], *ASXL1*-CHIP [OR=0.74], and *JAK2*-CHIP [OR=0.48] (**Fig 1C, middle; Supp Table 2**). Furthermore, rs80093687 carried a reduced risk of MyMs [OR=0.80, 95% CI=0.70-0.90], including myeloproliferative neoplasm (MPN) [OR=0.63, 95% CI=0.50-0.81] and acute myeloid leukemia (AML) [OR=0.83, 95% CI=0.72-0.95] (**Fig 1C, bottom**), and PheWAS strongly associated this variant with reduced peripheral counts of several blood cell lineages (**Fig 1D**)^11^. Importantly, no other phenotypic associations were identified for this variant, suggesting a selective impact on hematopoiesis (**Supp Fig 1E**). These associations suggested that the CHIP resilience haplotype at the 17q22 locus might selectively alter hematopoiesis to confer a pan-protective effect against CHIP driver mutations and MyM.

### rs17834140-T protects from CHIP through HSC-selective downregulation of MSI2

To investigate mechanisms underlying inherited CHIP-resilience, we conducted fine mapping at the 17q22 haplotype (**Supp Table 3**). While no candidate variants mapped to coding regions or were predicted to alter splicing, we identified rs17834140 – a statistically fine-mapped SNP in perfect LD with rs80093687 [R^2^=1.0] – as a likely causal variant that was compelling for several reasons. First, rs17834140-T resided within a putative regulatory element bound by a complex of transcription factors critical for maintaining human HSCs (**Fig 2A**)^27,28^. Second, the element was selectively accessible in HSCs (**Fig 2B**), but not in more differentiated progenitors or mature blood cells, nor in hematopoietic cell lines (**Supp Fig 2A**), suggesting a stem cell specific regulatory function. Third, H3K27ac-HiChIP in human HSPCs^29^ showed the element could physically contact the promoter of its proximal gene, *MSI2* (**Fig 2C**) – a potent RNA-binding regulator of a variety of tissue stem cells^30,31^, which has been implicated in aggressive cancers^32^. Importantly, *MSI2* was the only gene appreciably expressed within 1 Mb of rs17834140 in human HSCs (**Supp Fig 2B**), and there was high correlation between variant accessibility and *MSI2* expression across human hematopoietic cells [ATAC-RNA r=0.65] (**Supp Fig 2C**). Furthermore, CRISPR interference with dCas9-KRAB mediated repression of the regulatory element reduced *MSI2* expression in CD34^+^CD45RA^-^CD90^+^ HSC-enriched populations (**Fig 2D**). These insights suggested that the rs17834140-harboring regulatory element acts as an endogenous enhancer of *MSI2*.

**Figure 2:**
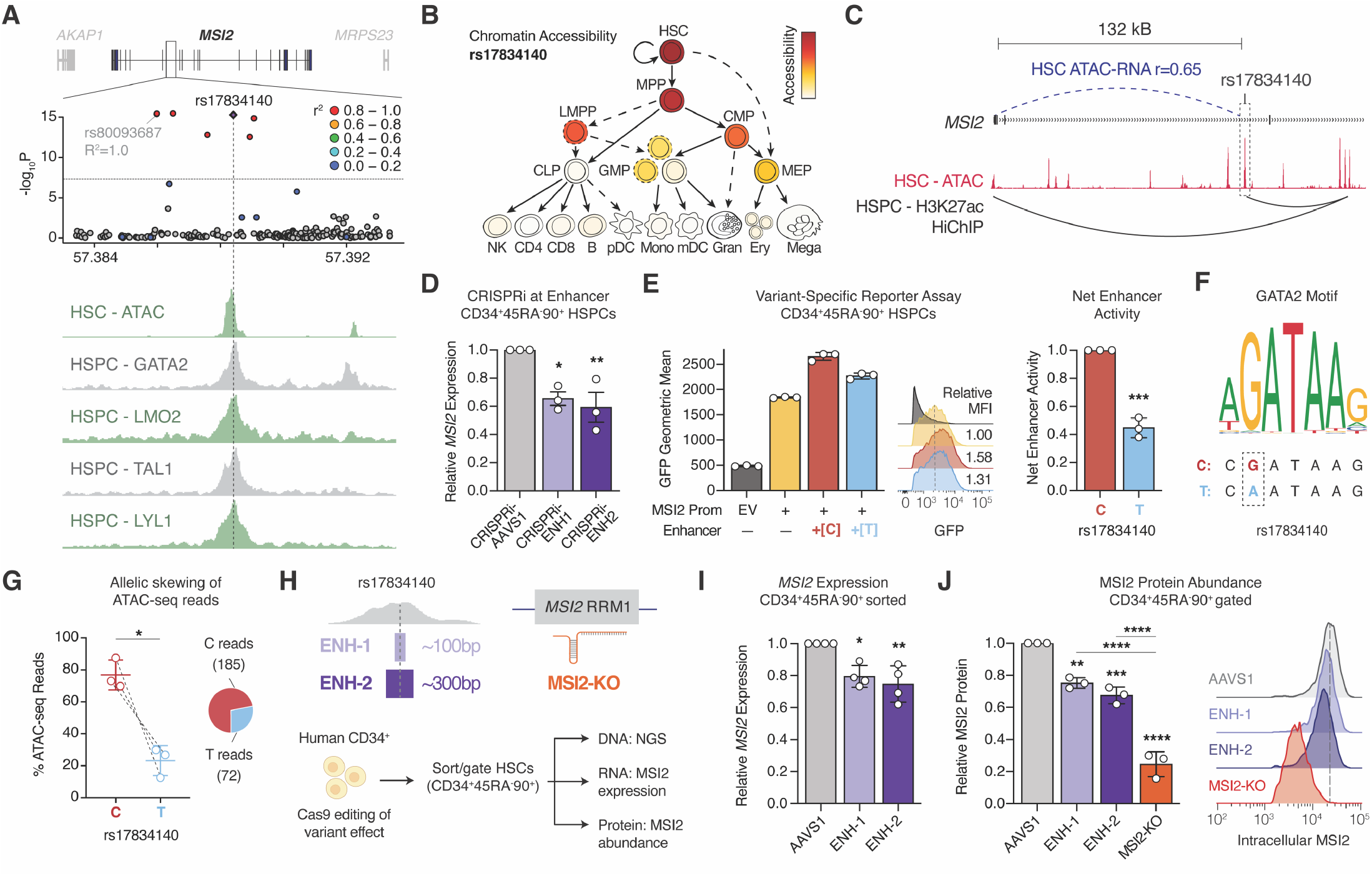
Regulatory variant rs17834140-T protects from CHIP through HSC-selective downregulation of *MSI2*. **(A)** LocusZoom plot identifying rs17834140 at the 17q22 CHIP-resilience haplotype (R^2^=1.0 with rs80093687) as a putative causal regulatory variant, with tracks for chromatin accessibility (ATAC) and transcription factor binding (ChIP-seq) in primary human HSPCs. **(B)** Normalized chromatin accessibility of the rs17834140-harboring regulatory element in 19 hematopoietic cell types, indicating HSC-selective regulatory activity. **(C)** H3K27ac-HiChIP track showing physical contact between rs17834140 and the *MSI2* promoter in human HSPCs^29^. **(D)** Reduced *MSI2* mRNA expression in CD34^+^CD45RA^-^CD90^+^ HSC-enriched cells, 3 days after CRISPR interference targeting the rs17834140-harboring regulatory element. *N*=3 donors. **(E)** Reporter assay showing enhancer activity of rs17834140 wild-type (C) or CHIP-resilience (T) alleles in CD34^+^CD45RA^-^CD90^+^ HSC-enriched cells. *N*=3 donors. **(F)** rs17834140 is predicted to disrupt a conserved GATA2 binding motif, a factor known to bind at this site. **(G)** Allele-specific ATAC-sequencing reads at rs17834140 in HSPCs showing reduced chromatin accessibility with the resilience (T) allele. *N*=3 donors. **(H)** Editing strategy modeling rs17834140-T effect with ∼100bp (ENH-1) or ∼300bp (ENH-2) microdeletions, or MSI2-knockout (MSI2-KO). **(I)** Reduced *MSI2* mRNA expression in CD34^+^CD45RA^-^CD90^+^ HSC-enriched cells, 3 days after enhancer microdeletions. *N*=4 donors. **(J)** Flow cytometry showing reduced intracellular MSI2 protein abundance in CD34^+^CD45RA^-^CD90^+^ gated HSC-enriched cells, 3 days after editing. *N*=3 donors.

To decipher variant effects, we introduced the rs17834140-harboring regulatory element upstream of a *MSI2*-promoter driven GFP reporter in primary human HSPCs (**Supp Fig 2D**). Within transduced CD34^+^CD45RA^-^ CD90^+^ HSC-enriched cells, addition of the wild-type (rs17834140-C) element augmented reporter expression, validating that this element acts as an enhancer in HSCs (**Fig 2E**). Remarkably, mutagenesis to recreate the CHIP-resilience variant (rs17834140-T) alone in otherwise matched constructs reduced its enhancer activity by ∼2-fold. We examined canonical motifs at this element, observing that rs17834140-T was predicted to break a highly conserved GATA-binding site (CADD score: 15.6) (**Fig 2F, Supp Fig 2E**), which was notable given that GATA2 is known to bind strongly at this enhancer (**Fig 2A**). Consistent with this, we observed a depletion of T-alleles in ATAC-sequencing reads from HSPCs of heterozygote donors (**Fig 2G**). These data demonstrate that rs17834140-T causes loss-of-function at an *MSI2* enhancer in human HSCs.

We next sought to faithfully model resilience variant effects in primary CD34^+^ human HSPCs. We utilized CRISPR/Cas9 to minimally excise the putative GATA2-binding site with ∼100bp (ENH-1) or ∼300bp (ENH-2) microdeletions harboring rs17834140 (**Fig 2H**). We bench-marked phenotypes against edits targeting the RNA-recognition motif (RRM)-1 domain of MSI2 (KO). Three days after editing, we observed a high efficiency of microdeletions (∼70-80%) and MSI2-KO (∼90%) (**Supp Fig 2F-I**). Regulatory element microdeletions reduced *MSI2* mRNA expression in CD34^+^CD45RA^-^CD90^+^ HSPCs relative to safe-harbor *AAVS1*-edited controls (**Fig 2I**), selectively within molecularly defined HSCs (**Supp Fig 2J**) with no impact on other genes at the locus (**Supp Fig 2K**). We additionally performed intracellular flow cytometry, finding concordant decreases in MSI2 protein abundance in CD34^+^CD45RA^-^ CD90^+^ HSPCs following ENH-1 (by ∼20% vs. AAVS1) and ENH-2 (by ∼30% vs. AAVS1) microdeletions (**Fig 2J**). Thus, our functional analyses indicate that rs17834140-T protects from CHIP through HSC-selective downregulation of MSI2 levels.

### Genetic variation-driven loss of MSI2 enhancer reduces human HSC fitness

We next sought to understand how inherited CHIP resilience impacts human HSCs (**Fig 3A**). Six days after editing, MSI2-KO did not impact the proportion of CD34^+^CD45RA^-^ cells in culture, but reduced phenotypic long-term (LT-) HSCs (CD34^+^CD45RA^-^CD90^+^CD133^+^EPCR^+^ITGA3^+^) principally through loss of primitive CD90^+^ HSPCs (**Fig 3B**), consistent with the role of MSI2 in HSC maintenance^30–33^. Remarkably, ENH-1 and ENH-2 microdeletions of the rs17834140-site were sufficient to partially phenocopy this effect, both in proportion (**Fig 3C**) and total number (**Fig 3D, Supp Fig 3A**) of LT-HSCs. In an orthogonal model, we employed recently developed cytokine-free and chemically defined cultures of human cord blood CD34^+^ HSCs^34^. In these conditions, after 14 days, we further observed loss of LT-HSC maintenance (**Fig 3E**), with total LT-HSC numbers significantly reduced following ENH-1 (by 44% vs. AAVS1) or MSI2-KO (by 85% vs. AAVS1) editing (**Fig 3F, Supp Fig 3B**). Collectively, these data suggest that loss-of-function at the *MSI2* enhancer through genetic variation might alter LT-HSC maintenance.

**Figure 3:**
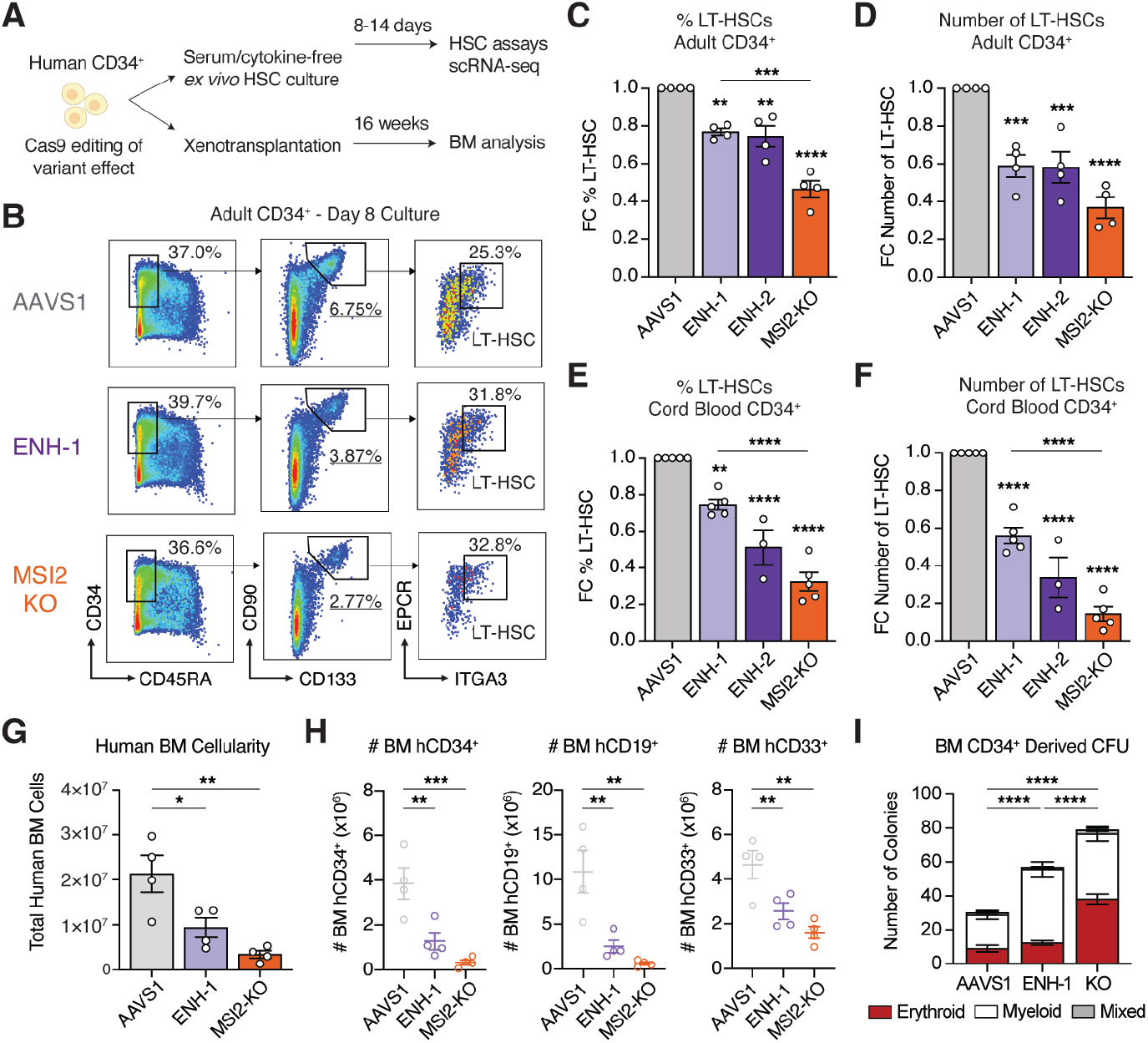
Genetic variation-driven loss of MSI2 enhancer reduces human HSC fitness. **(A)** Outline of HSC assays used to assess CHIP-resilience variant effect. **(B)** Representative flow cytometry gating for LT-HSC (CD34^+^CD45RA^-^CD90^+^CD133^+^EPCR^+^ITGA3^+^) immunophenotyping, six days after editing adult CD34^+^ HSPCs. **(C-D)** Proportion and total numbers of phenotypic LT-HSCs on day 8 of serum-free culture (6 days post-editing) of adult CD34^+^ HSPCs, relative to AAVS1-edited controls. *N*=4 donors. **(E-F)** Proportion and total numbers of phenotypic LT-HSCs on day 14 of cytokine-free culture (11 days post-editing) of cord blood CD34^+^ HSPCs, relative to AAVS1-edited controls. *N*=3-5 donors. **(G)** Total human cells achieving long-term engraftment in bone marrow (BM) of NBSGW mice, 16-weeks after transplantation with AAVS1, ENH-1 or MSI2-KO edited cord blood HSPCs. *N*=12 mice. **(H)** Total human CD45^+^ engrafted cells in BM expressing lineage markers (CD34^+^ HSPCs, CD19^+^ B-cells, CD33^+^ myeloid cells). *N*=12 mice. **(I)** Number of colonies formed by BM-selected human CD34^+^ cells cultured in methyl-cellulose, 16 weeks post-transplantation. *N*=12 mice.

To determine if the rs17834140-T-harboring regulatory element could modulate *in vivo* self-renewal and HSC function, we performed xenotransplantation experiments involving NBSGW (Kit-mutant and immunodeficient NOD.Cg-Kit^W41J^Tyr^+^ Prkdc^scid^Il2rg^tm1Wjl^/ThomJ) mice^35,36^. Mice transplanted with ENH-1 edited human cord blood-derived CD34^+^ cells had reduced total human bone marrow (BM) cellularity (**Fig 3G, Supp Fig 3C**) and multilineage engraftment of CD34^+^ HSPCs, CD19^+^ B-cells, and CD33^+^ myeloid cells (**Fig 3H**) at 16 weeks post transplantation, partially phenocopying a more severe defect in BM reconstitution observed with MSI2-KO edited cells. Among engrafted human CD45^+^ cells, ENH-1 or MSI2-KO edited cells had reduced proportions of CD34^+^ HSPCs and CD19^+^ B-cells in BM and spleen (**Supp Fig 3D**), representing a loss of HSCs with multilineage potential. Interestingly, there was a greater percentage of CD33^+^ myeloid cells and myeloid colony formation for human CD34^+^ cells (**Fig 3I, Supp Fig 3E**), suggestive of increased myeloid lineage commitment. Consistent with the role of MSI2 in promoting symmetric self-renewal over asymmetric and differentiative HSC divisions^32,37^, these data support the idea that CHIP-resilient HSCs have reduced long-term functional maintenance *in vivo*, through a greater propensity for differentiation at the expense of self-renewal capacity.

### Modified stem cell RNA network protecting HSCs from CHIP and MyM

Our functional studies led us to investigate mechanisms by which genetic variation-driven downregulation of MSI2 reduces HSC fitness and protects from CHIP. As MSI2 is an RNA-binding protein that regulates stability and translation^38^ of mRNAs, we sought to define its direct binding targets. Given that prior studies were conducted in cell lines^33^ or mouse cells^38^, the MSI2-regulated network in human HSCs has remained undefined. We therefore utilized MSI2-HyperTRIBE^38^ (**Fig 4A**) – in which a fusion MSI2-ADAR (RNA-editing enzyme) protein catalyzes A>G edits on MSI2-bound RNA targets – in human cord blood-derived CD34^+^ HSPCs. MSI2-HyperTRIBE identified 12,928 edits across 3,614 genes with between 1-62 editing sites per gene (**Supp Fig 4A-C, Supp Table 4**). MSI2-bound mRNAs were enriched for the UAG motif in their 3’ UTRs (**Fig 4B**), consistent with prior reports that identified a more limited set of targets^33,38^, and there was strong concordance in bound mRNAs with orthogonal CLIP-seq targets of MSI2^33^ (**Supp Fig 4D-E**).

**Figure 4:**
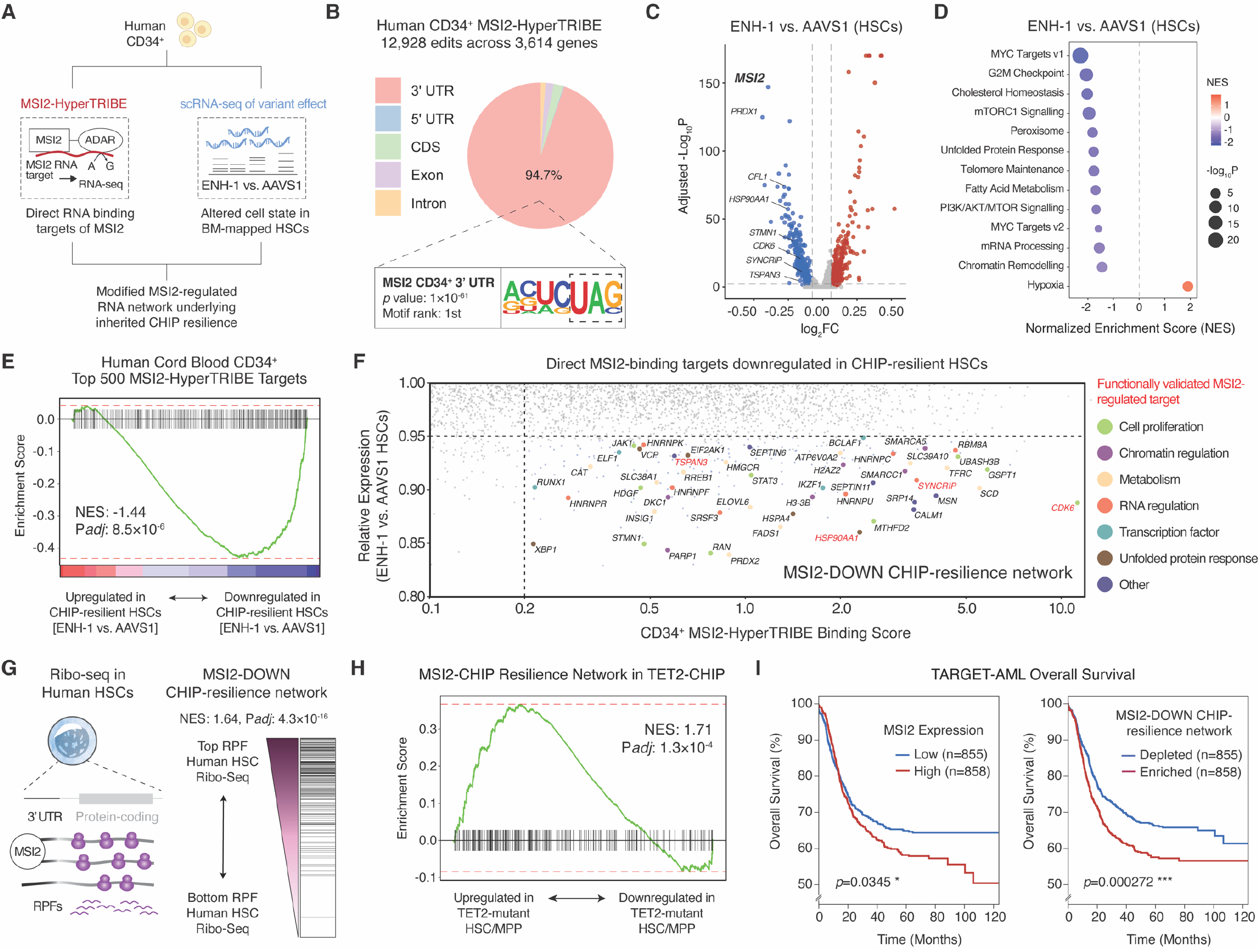
Modified RNA regulation underlying CHIP-resilience in human HSCs. **(A)** Overview of approach to determine MSI2-regulated RNA network in human HSCs. **(B)** MSI2-HyperTRIBE edit sites are enriched in 3’ UTR of target mRNAs with the UAG MSI2-recognition motif. **(C)** Volcano plot of differentially expressed genes (DEGs) between ENH-1 vs. AAVS1-edited HSCs. **(D)** Gene set enrichment analysis (GSEA) of ENH-1 vs. AAVS1 edited HSCs. **(E)** CHIP-resilient HSCs (ENH-1 vs. AAVS1) show negative enrichment for CD34^+^ MSI2-HyperTRIBE mRNA targets. **(F)** Identification of an RNA network that is MSI2-bound and functionally downregulated in CHIP-resilient HSCs. **(G)** Ribo-seq profiling of human HSCs with GSEA showing CHIP-resilience network mRNAs are enriched for ribosome protected fragments (RPFs). **(H)** *TET2*-mutant HSC/MPPs show positive enrichment for the MSI2-DOWN CHIP-resilience network, relative to their wild-type counterparts from the same donors. **(I)** Overall survival of pediatric acute myeloid leukemia (AML) patients in the TARGET-AML cohort, stratified by expression of either *MSI2* or the MSI2-DOWN CHIP-resilience network. Log-rank test.

We then investigated how reduced MSI2 levels impact its network to confer HSC resilience to CHIP. scRNA-seq was performed in human CD34^+^CD45RA^-^CD90^+^ HSC-enriched cells with AAVS1, ENH-1, or MSI2-KO edits and analyses were focused on molecularly defined HSCs (**Supp Fig 4F-G**). CHIP-resilient HSCs (ENH-1 vs. AAVS1) had 24% loss of MSI2 expression and 1,005 differentially expressed genes (DEGs) (**Fig 4C, Supp Table 5**) with high concordance to the MSI2-knockout signature [Pearson’s r=0.75] (**Supp Fig 4H**), representing pathways including downregulation of proliferation, MYC targets, and cholesterol biosynthesis genes (**Fig 4D**). Strikingly, there was strong negative enrichment of MSI2-HyperTRIBE targets in ENH-1 microdeleted HSCs with reduced MSI2 levels, suggesting that most MSI2-bound mRNA targets were downregulated in CHIP-resilient HSCs (**Fig 4E**).

To understand direct MSI2 bound targets that are functionally altered in human HSCs, we next overlapped MSI2-HyperTRIBE and scRNA-seq datasets. More highly edited MSI2 binding targets (mRNAs with ≥20% editing by MSI2-ADAR) and significantly downregulated genes were selected. This identified a network of 208 direct MSI2-binding targets that were also downregulated in ENH-1 vs. AAVS1 HSCs, hereafter termed MSI2-DOWN genes (**Fig 4F, Supp Table 6**). This network was enriched for functional MSI2-targets (e.g., *TSPAN3*^39^, *HSP90AA1*^33^, *SYNCRIP*^40^, *CDK6*^41^), validating our approach, as well as positive regulators of human HSCs (e.g., *TFRC*^42,43^, *DKC1*^44^, *H3-3B*^45^), but with no previously studied role in CHIP, nominating candidate regulators of stem cell clonality downstream of MSI2. We leveraged ribosome profiling in primary human CD34^+^CD45RA^-^CD90^+^ HSCs^36^, finding that the MSI2-DOWN CHIP-resilience network genes were enriched for higher numbers of ribosome protected fragments (RPFs) in comparison to other transcripts (**Fig 4G**). This indicated that the RNA network positively regulated by MSI2 are among the more highly translated mRNAs in human HSCs, validating the functionality of this network of RNAs that is modulated through genetic variation.

Having identified an RNA network underlying CHIP resilience in HSCs, we sought to understand its clinical relevance. We leveraged TARGET-seq (joint single-cell genotyping and RNA-sequencing) data of CHIP mutant vs. wild-type HSCs from the same donors, enabling insight into the transcriptional impact of CHIP mutations in an isogenic setting^46^. Strikingly, the MSI2-DOWN network (downregulated in CHIP-resilient HSCs) was conversely upregulated in TET2 mutant HSCs in comparison to their wild-type counterparts (**Fig 4H**). To explore the prognostic value of the MSI2-DOWN network in MyMs, we interrogated the TARGET AML cohort^47^. Patients with an enrichment of either MSI2 expression itself or of its downstream network displayed a significantly worse prognosis (**Fig 4I**). Collectively, we show that the MSI2-DOWN CHIP-resilience network is enriched for positive regulators of HSC proliferation, that may promote stem cell clonal advantage in CHIP and MyMs, suggesting that inherited downregulation of this network might neutralize the impact of acquired CHIP mutations.

### MSI2 levels modify clonal dominance of ASXL1-mutant HSCs

Finally, we sought to understand how altered MSI2 levels in human HSCs impact progression of CHIP. We studied the effect of the protective rs17834140-T variant on clonal dynamics of CHIP in a cohort with targeted sequencing of two serial blood samples taken ∼6 years apart (**Fig 5A**). Among 513 individuals with persistent CHIP (second VAF also ≥2%), rs17834140-T carriers had a significantly slower measured growth rate of mutant CHIP clones, with many clones showing regression or minimal growth over ∼6 years of sampling (median growth rates: C/C = 0.12%/year, C/T = - 0.31%/year) (**Fig 5B**). Indeed, rs17834140-T carriers had greater odds of their CHIP mutation being transient (second VAF <2%) [OR=1.82, 95% CI: 1.29-2.52, p=0.0007] (**Fig 5C**), which was most pronounced for ASXL1-CHIP [OR=2.72, 95% CI: 1.00-7.33, p=0.057]. Together, these data provide evidence that rs17834140-T attenuates CHIP progression through longitudinal targeted sequencing in humans.

**Figure 5:**
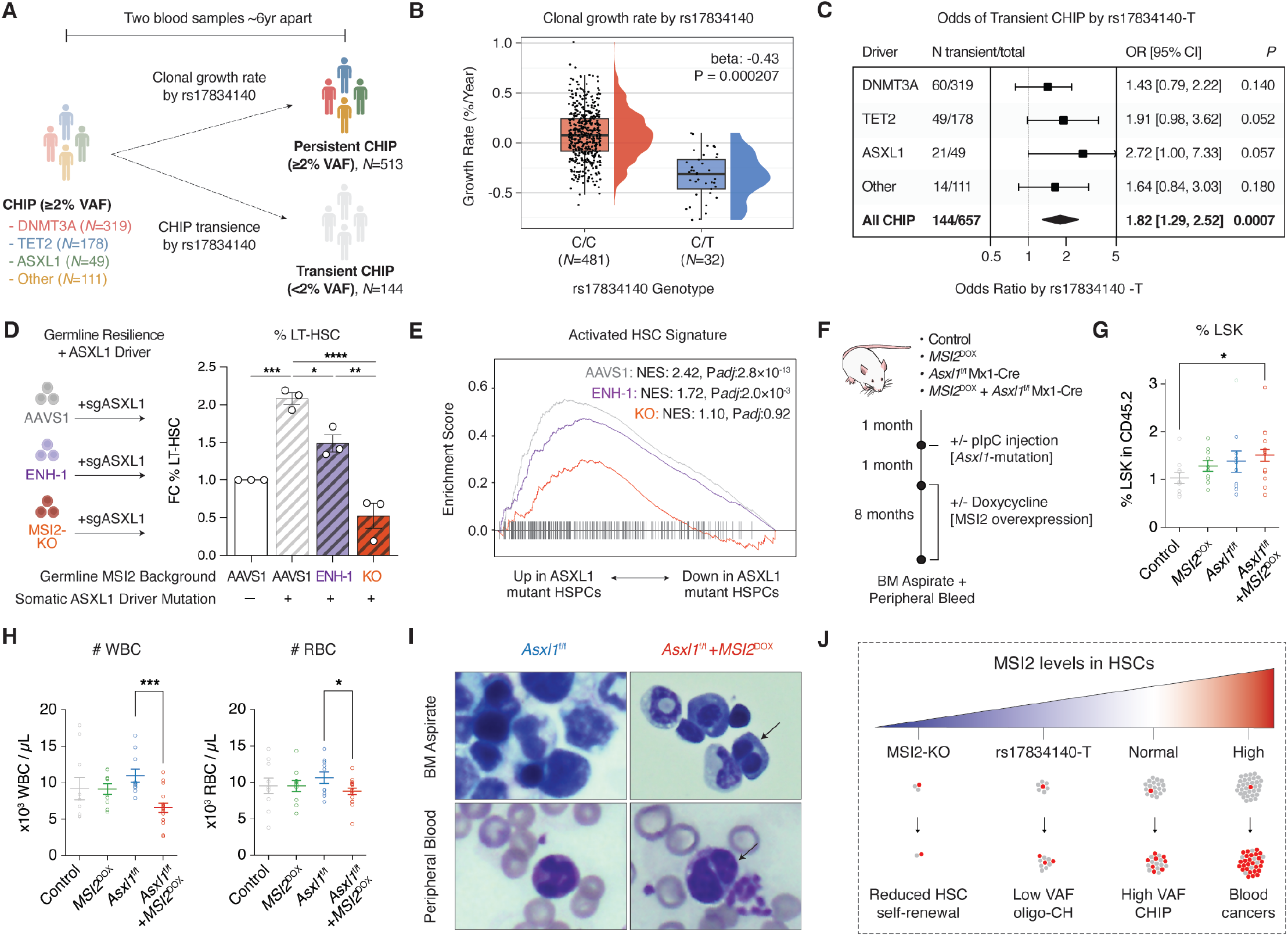
Stem cell MSI2 levels modify clonal dominance of *ASXL1*-mutant HSCs. **(A)** Approach to assess associations between rs17834140-T and longitudinal CHIP progression in a cohort with serial blood samples ∼6 years apart. **(B)** Growth rate [% VAF/year] of CHIP-mutations stratified by C/C (*N*=481) or C/T (*N*=32) genotype at rs17834140. **(C)** Odds of CHIP transience (>2% VAF at baseline, <2% VAF at follow-up) by rs17834140-T for driver mutations. **(D)** Dual modeling of germline CHIP-resilience (ENH-1 microdeletion, MSI2-knockout, AAVS1 control) with ASXL1-editing in primary adult CD34^+^ HSPCs, quantifying LT-HSCs 7 days post-editing. *N*=3 donors. **(E)** RNA-seq of CD34^+^CD45RA^-^CD90^+^ HSC-enriched cells showing that ASXL1 mutations enrich for an activated HSC signature^42^, which is attenuated by MSI2 loss (ENH-1 microdeletion or MSI2-knockout). **(F)** Workflow of murine model evaluating MSI2 and Asxl1 cooperativity. **(G)** Quantification of Lin^-^ Sca-1^+^c-Kit^+^ (LSK) cells, 1-month post-doxycycline treatment in murine model. *N*=10-17 mice per condition. **(H)** Peripheral red blood cell (RBC) and white blood cell (WBC) counts 8-months post-doxycycline in murine model. *N*=10-16 mice per condition. **(I)** Representative blood smears from bone marrow and peripheral blood, showing myelodysplastic syndrome (MDS) features (hyposegmented Pelger-Huët-like neutrophils and binucleate erythroid precursors) in *Asxl1*^f/f^+*MSI2*^DOX^ mice. **(J)** Model of how MSI2 levels in HSCs regulate stem cell pool size (grey) and clonal advantage of somatic CHIP driver mutations (red).

To validate MSI2 levels as a modifier of CHIP, we focused on ASXL1-mutant CHIP, a common driver in 20% of MDS/AML, and for which rs17834140-T carried amongst the most protective associations for CHIP prevalence [OR=0.77, 95% CI: 0.66-0.90] and transience. We modeled patient *ASXL1* mutations (**Supp Fig 5A**) in primary human CD34^+^ HSPCs with an exon 12-targeting guide RNA that increases maintenance of phenotypic LT-HSCs in culture (**Supp Fig 5B**) and recapitulates clonal expansion in a xenograft model^48^. HSPCs were edited to model germline CHIP resilience (80-100% editing) and acquired *ASXL1*-mutations (91-95% editing), with most alleles co-edited at both loci (**Supp Fig 5C**). While *ASXL1*-editing expanded the proportion of LT-HSCs present in ex vivo cultures, this was partially or completely attenuated with a background of ENH-1 micro-deletion or MSI2-KO (**Fig 5D**). RNA-sequencing within sorted CD34^+^CD45RA^-^CD90^+^ cells indicated that *ASXL1*-edited HSPCs were enriched for a more activated HSC signature^42^, which was reversed with concurrent ENH-1 microdeletion or MSI2-KO (**Fig 5E**), among other transcriptional pathways (**Supp Fig 5D**). These data suggest that inherited CHIP-resilience through MSI2 downregulation can protect HSCs from expansion mediated by *ASXL1*-mutations.

Conversely, stem cell MSI2 levels are commonly upregulated in MyM, and drive aggressive disease with poor prognosis. To probe the interaction of MSI2 levels with CHIP in this disease setting, we generated a mouse model to probe the synergistic impact of MSI2-overexpression and *Asxl1*-mutant CHIP (**Fig 5F, Supp Fig 5E**). While MSI2-overexpression or *Asxl1*-deletion alone had minimal impact on the frequency of HSC-enriched Lin^-^Sca-1^+^c-Kit^+^ (LSK) cells, mice transplanted with *Asxl1*^f/f^+*MSI2*^DOX^ expressing cells had a 1.5-fold average increase in LSK cell frequency (**Fig 5G**). Over 8 months, while *Asxl1*^f/f^ and *MSI2*^DOX^ mice did not show evidence of hematologic disease, *Asxl1*^f/f^+*MSI2*^DOX^ mice developed significantly reduced peripheral leukocyte and red blood cell counts (**Fig 5H**), along with other blood trait alterations (**Supp Fig 5F**), and blood smears showed MDS features that included hyposegmented Pelger-Huët-like neutrophils and binucleate erythroid precursors (**Fig 5I, Supp Fig 5G**). Taken together, our data suggest a model in which MSI2 levels in HSCs functionally interact with CHIP/MyM driver mutations by modifying their clonal advantage both prior to and after somatic mutation acquisition (**Fig 5J**).

## Discussion

Genetic variation provides opportunities to uncover natural mechanisms for disease resilience. Notable examples include loss-of-function (LoF) variants in *CCR5* conferring HIV resistance^49^, *PCSK9* protecting from cardiovascular disease^50^, and *BCL11A* suppression enabling amelioration of sickle cell disease and beta-thalassemia^51,52^ – insights that have been harnessed for therapeutic development. Despite tremendous potential – and while much has been learned about cancer predisposition – the study of inherited cancer resilience is largely unexplored. As purifying selection and sample sizes hinder detection of rare LoF variants^53^, we reasoned that large population biobanks might enable such common protective variation to be identified. In this study, we identify a protective mechanism against CHIP – an exemplar of somatic mosaicism that serves as a precursor to MyM – through genetic downregulation of the expression and function of the RNA-binding protein MSI2 in HSCs.

Our studies provide critical insight into RNA networks underlying HSC maintenance and selective advantage in CHIP. In recent years, RNA-binding proteins have emerged as key regulators of the post-transcriptional circuitry that play a central role in self-renewal and differentiation in stem cells^54^. While previous work established MSI2 as a potent regulator of HSC numbers *in vivo*^32,33,37^, its downstream mechanisms and the fact that it is impacted by naturally occurring human genetic variation has remained unknown. We identified, for the first time, the direct binding targets of MSI2 in human HSCs, revealing a network enriched for positive regulators of HSC maintenance. Importantly, MSI2-regulated mRNAs are upregulated in HSCs with acquired CHIP driver mutations^46^. Our study exemplifies how genetic variation may downregulate this network to enable CHIP resilience. Systematic perturbation of this network in future studies is likely to deepen our understanding of the mechanisms involved in HSC maintenance and the fitness advantage in CHIP. This may also be of broader relevance in MSI2-driven lymphoid malignancies^41,55,56^ and solid tumors^57–60^.

Our data provide human genetic evidence supporting MSI2 inhibition as a potential therapeutic strategy for CHIP or in those at high risk for acquiring MyM for other reasons. We show that partial and selective downregulation of MSI2 in HSCs is adaptive through conferring CHIP/MyM resilience (**Fig 5J**), without other apparent adverse implications in variant-carriers. Conversely, MSI2 is frequently overexpressed in AML^61,62^, and upregulated during CML progression^62^, and we show that MSI2 overexpression cooperates with Asxl1-mutations to induce a MyM in a mouse model. Together, our results support two (non-mutually exclusive) models, in which reduced MSI2 levels may: i) reduce the baseline HSC pool (lowering the likelihood of any cell acquiring a CHIP driver mutation); ii) attenuate clonal advantage of drivers once acquired. Our study highlights the potential to target MSI2, through small molecule inhibition or genome editing at its enhancer, for blood cancer prevention. More broadly, we provide an example of how resilience to cancer can arise through inherited genetic variation, motivating the search for other natural pathways that could be leveraged to prevent or treat malignancy.

## Acknowledgements

We are grateful to members of the Sankaran Lab for valuable comments and suggestions. This work was supported by the Howard Hughes Medical Institute (V.G.S.), the Mather’s Foundation (V.G.S.), the Bill and Melinda Gates Foundation (V.G.S.), and National Institutes of Health (NIH) grants R01 DK103794, R01 CA265726, R01 CA292941, R33 CA278393, and R01 HL146500 (V.G.S.). A-L.N. is supported by the EMBO Postdoctoral Fellowship (ALTF 209-2024). A.G.B. is supported by NIH grants DP5 OD029586, R01 AG088657, R01 AG083736, a Burroughs Wellcome Fund Career Award, a Pew-Stewart Scholar for Cancer Research award and a Hevolution/AFAR New Investigator Award in Aging Biology and Geroscience Research. V.G.S. is an Investigator of the Howard Hughes Medical Institute.

## Author contributions

G.A., M.G.K., and V.G.S. conceptualized the study. G.A., M.G.K. and V.G.S. devised the methodology. G.A., M.A., X.X., P.L., T-C.H., T.S.B., A.C., A.S., and J.T. performed experiments. G.A., Y.P., C-L.P., A.J.L., L.N., D.R., and K.L.M. performed computational analyses. C.G., T.Y., A-L.N., L.d.V., L.W., B.H.D., O.A-W., J.M.A., R.S., and A.G.B. provided resources and feedback. G.A., Y.P., U.P.A., A.J.L., M.G.K., and V.G.S. provided data visualization media.

V.G.S. acquired funding for this work and provided overall project oversight. G.A. and V.G.S. wrote the original manuscript, as well as edited the manuscript with input from all authors.

## Data availability

Raw and processed data have been deposited at GEO, and will be available at the time of publication.

## Materials and Methods

### All of Us and GWAS meta-analysis

All of Us (AoU) is a longitudinal cohort containing short-read whole genome sequencing data (mean depth of 30x). Informed consent is in place for all AoU participants, and the protocol was reviewed by the Institutional Review Board (IRB) of the AoU Research Program. Variant-level QC metrics included QUAL > 60, ExcessHet < 54.69, GQ > 20, DP > 10, AB > 0.2 for heterozygotes. Variants with population-specific allele frequency (AF) > 1% or population-specific allele count (AC) > 100 were selected. Sites with more than 100 distinct alternate alleles were excluded.

Individual-level CHIP calls in AoU were ascertained, as previously described^63^. Only samples with matching reported sex and genetically inferred sex were included, and analyses were performed on individuals of European ancestry with no diagnosis of hematological malignancy (using filtering criteria as previously reported^63^). PLINK (v1.9) was used to perform logistic regression with Firth correction to assess associations with CHIP for the most significant sentinel SNP at 24 loci previously reported^11^. An inverse variance-weighted fixed-effect meta-analysis was performed across AoU associations and summary statistics in UK Biobank (UKB) and Geisinger Health Study (GHS) cohorts^11^. Additionally, the effect of rs80093687 by specific CHIP driver mutations (*DNMT3A, TET2, ASXL1, JAK2*) was evaluated as a meta-analysis (across AoU and UKB). All analyses in AoU were adjusted for age, sex at birth and the first 5 genetic principal components as covariates.

For associations with myeloid malignancy (MyM) risk, effect estimates for rs80093687 were extracted from GWAS summary statistics for myeloproliferative neoplasm (MPN) [3,797 cases and 1,152,977 controls^10^], acute myeloid leukemia (AML) [4,018 cases and 10,488 controls^64^] and myelodysplastic syndromes (MDS) [907 cases and 5,604 controls^65^]. For phenome-wide association study (PheWAS), previously reported summary statistics for rs80093687^11^ were plotted using the PheWAS package in R.

### Fine-mapping and variant annotation

Statistical fine-mapping was performed using GWAS summary statistics for CHIP in the UKB European ancestry cohort^11^ [GWAS catalogue: GCST90165267], and linkage disequilibrium (LD) matrices for UKB participants of British ancestry^66^. Summary statistics were converted from hg38 to hg19 using LiftOver. Variants with minor allele frequency (MAF) > 0.001 within 1.5 Mb of the sentinel SNP rs80093687 (7,012 in total) were fine-mapped using SuSIE^67^, CARMA^68^ and FINEMAP^69^, allowing up to 10 causal variants and reporting 95% credible sets for each method. All methods identified the same six variants (rs80093687, rs188761458, rs199691861, rs118121072, rs17834140, rs150497606) in a single 95% credible set.

Fine-mapped variants and those in strong LD with rs80093687 (R^2^ ≥ 0.8, ± 1.5 Mb) were annotated with genomic context, splicing predictions (using SpliceAI^70^), and chromatin accessibility across 13 hematopoietic cell types^71^. Normalized mRNA expression for all genes within 1 Mb of rs17834140 was extracted from a single-cell multiome dataset of human HSCs (old-1 and old-2 donors^4^). For ATAC-RNA correlation, hematopoietic cell types were clustered into N=75 ‘metacells’ to address single-cell sparsity, and Pearson’s correlation was computed between mean *MSI2* expression and chromatin accessibility at the rs17834140-harboring regulatory element. H3K27ac-HiChIP and ChIP-seq data (GATA2, LMO2, TAL1, LYL1) in CD34^+^CD45RA^-^ HSPCs were previously reported^29^. Transcription factor motif analysis was conducted using motifBreakR (v2.16)^72^, with HOCOMOCO v11 matrices and a statistical threshold of 1×10^-3^. Allelic skewing in ATAC-seq data of primary human CD34^+^ HSPCs was assessed in three heterozygotes for rs17834140 (GEO accessions GSE194122, GSE219015 and GSE156733). Raw FASTQ files were aligned to hg38 using bwa (v0.7.18), BAM files were generated with samtools (v1.20), and bcftools mpileup was used to call ‘C’ and ‘T’ alleles at rs17834140.

### Primary cell culture

Primary human CD34^+^ HSPCs from mobilized peripheral blood of healthy donors were obtained from the Fred Hutchinson Cancer Research Center. Thawed cells were cultured at 5×10^5^ cells/mL in serum-free StemSpan SFEM II medium (StemCell Technologies) supplemented with 1% L-glutamine (ThermoFisher Scientific, 25-030-081), 1X penicillin/streptomycin (Life Technologies, 15140-122), 1X CC100 (containing the cytokines FLT3L, SCF, IL-3 and IL-6; StemCell Technologies, 02690), 100 ng/mL recombinant thrombopoietin (TPO; PeproTech, 300-18), and 35 nM UM171 (StemCell Technologies).

CD34^+^ HSPCs from cord blood (CB) of healthy newborns (Pasquarello Tissue Bank, Dana-Farber Cancer Institute) were purified using the EasySep Human Cord Blood CD34^+^ positive selection kit (StemCell Technologies) according to the manufacturer’s instructions, and cultured in cytokine-free conditions, as described^34^. In brief, CB HSPCs were cultured at 7×10^4^ to 1×10^5^ cells/mL in Iscove’s Modified Dulbecco’s Medium (IMDM; Life Technologies), supplemented with 1% L-glutamine (Thermo Fisher Scientific), 1% penicillin/streptomycin (Life Technologies), 1% insulintransferrin-selenium-ethanolamine (ITSX; Life Technologies), 1 mg/ml polyvinyl alcohol (PVA; Sigma-Aldrich), 1 µM 740Y-P (MedChemExpress), 0.1 µM butyzamide (Med-ChemExpress), and 70 nM UM171.

### CRISPR interference

For CRISPR interference, dCas9-KRAB (from Addgene plasmid #220838) was subcloned into the backbone of Addgene plasmid #204472, and in vitro transcription was performed to make purified mRNA encoding dCas9-KRAB. On day 2 of culture, HSPCs were washed two times in DPBS, and resuspended in 20 µL of Lonza P3 buffer with supplement. For each nucleofection, HSPCs were mixed with 2 µg dCas9-KRAB mRNA and total 2 µL sgRNA targeting either the *AAVS1* safe-harbor locus or the rs17834140-harboring regulatory element with independent pairs of sgRNAs [*AAVS1*: 1 µL each of sgAAVS1_g1 and sgAAVS1_g2; CRISPRi-ENH-1: 1 µL each of sgENH-1_g1 and sgENH-1_g2; CRISPRi-ENH-2: 1 µL each of sgENH-2_g1 and sgENH-2_g2 sgRNAs]. HSPCs were electroporated in 20 µL Nucleocuvette strips using the Lonza 4D Nucleofector, with the DS-130 program. Immediately after electroporation, 80 µL of prewarmed media were added to the electroporation cuvette, which was placed in an incubator at 37 °C for 5 mins. Cells were then plated at a density of 5×10^5^ cells/mL in adequate complete media. 72 hrs after nucleofection, HSPCs were sorted to enrich for HSCs (DAPI^neg^CD34^+^CD45RA^-^CD90^+^). Subsequently, total RNA was extracted, cDNA was generated, and real-time quantitative PCR (RT-qPCR) was performed to detect MSI2 expression (as described below).

### Lentiviral packaging and quantification

Lentiviral reporter constructs (as previously described^73^) carrying the enhancer element with wild-type (C) and resilience (T) alleles of rs17834140 [hg38: chr17:57,388,104-57,388,632] were cloned upstream of the human *MSI2* promoter driving GFP reporter expression. The *MSI2* promoter element was synthesized as a g-block [hg38: chr17:57,256,155-57,256,741] (IDT Technologies). For lentiviral packaging, HEK-293T cells were cultured at 37 °C in DMEM (Life Technologies) supplemented with 10% FBS and 1% penicillin/streptomycin. Cells were plated into 15 cm^2^ plates and grown to ∼70% confluency on the day of transfection per lentiviral construct. For each plate, 15 µg of psPAX2 packaging plasmid, 7.5 µg of pMD2.G envelope plasmid and 15 µg of construct were added in the presence of Opti-MEM media (Gibco, 31985-062) and Lipofectamine 3000 Transfection Reagent (Invitrogen, L3000001). Viral supernatants were harvested twice at 48 and 72 hrs post-transfection, and concentrated by ultracentrifugation (24,000 rpm for 2 hrs at 4 °C) in the SW32 rotor of Beckman Coulter ultracentrifuge. After ultracentrifugation, the supernatant was decanted and viral pellets were resuspended in ice cold PBS with 0.1% BSA, and frozen at -80 °C till further use. Lentiviral constructs were titrated using the Lenti-X™ Provirus Quantitation Kit (Takara, 631239) in K562 cells (cultured at 37 °C in IMDM supplemented with 10% FBS and 1% penicillin/streptomycin), as per manufacturer’s instructions.

### Reporter assay

For lentiviral transduction, concentrated virus was added to CD34^+^ HSPCs on day 1 of culture at equal titres, in the presence of 8 µg/mL cyclosporin H (Sigma-Aldrich, SML1575). HSPCs were then spinfected at 2,000 rpm for 90 mins at 37 °C. 72 hrs after infection, HSPCs were harvested, washed twice with PBS, and incubated for 30 mins with different fluorescent-labelled antibodies: 1:40 dilution of anti-human CD34 PerCP-Cyanine5.5 (clone 561, BioLegend, #343612), 1:50 dilution of anti-human CD45RA AlexaFluor-488 (clone HI100, BioLegend, #304114) and 1:100 dilution of anti-human CD90 PE (clone 5E10, BioLegend, #328110). Immuno-phenotypic analyses were performed to quantify geometric mean fluorescence intensity (MFI) of GFP^+^ transduced cells within CD34^+^CD45RA^-^CD90^+^ gated HSC-enriched cells for each lentiviral construct. Analyses were performed on LSR-Fortessa (BD Biosciences). Data were analyzed using the FlowJo software.

### CRISPR/Cas9 RNP nucleofection and editing analysi

Electroporation was performed on either day 2 (for CD34^+^ from mobilized peripheral blood of healthy donors) or day 3 (for cord blood CD34^+^ from healthy newborns) of culture, using the Lonza 4D Nucleofector with 20 µL Nucleocuvette strips. The Cas9 ribonucleoprotein (RNP) complexes were prepared by combining 2.1 µL of Lonza P3 primary cell nucleofection reagent (Lonza, V4XP-3032), 1.7 µL of 62µM Alt-R S.p. HiFi Cas9 Nuclease V3 (IDT, 1081061) and total 1.0 µL of 100 µM sgRNA in IDTE pH 7.5 (IDT) [*AAVS1*: 0.5 µL each of sgAAVS1_g1 and sgAAVS1_g2; ENH-1 microdeletion: 0.5 µL each of sgENH-1_g1 and sgENH-1_g2; ENH-2 microdeletion: 0.5 µL each of sgENH-2_g1 and sgENH-2_g2; MSI2-KO: 1 µL of sgMSI2-KO]. HSPCs were washed two times in DPBS, then resuspended in 20 µL of Lonza P3 buffer with supplement (in presence of nucleofection enhancer). Cells were electroporated using the DZ-100 program in a 4D-Nucleofector X Unit (20 µL cuvettes). Immediately after electroporation, 80 µL of prewarmed media were added to the electroporation cuvette, which was placed in an incubator at 37 °C for 5 mins. Cells were then plated at a density of 5×10^5^ cells/mL in adequate complete media.

For analysis of DNA editing, CD34^+^CD45RA^-^CD90^+^ HSC-enriched cells were harvested for genomic DNA extraction at least 72 hrs post-nucleofection. PCR fragments flanking the editing site (at least 250 bp upstream and down-stream) were amplified and editing frequencies and outcomes were detected by either Sanger sequencing (and analyzed with the ICE analysis tool from Synthego) or next-generation sequencing (NGS). For NGS, Premium PCR Sequencing was performed by Plasmidsaurus using Oxford Nanopore Technology with custom analysis and annotation, and analysis was performed using CRISPResso2^74^.

### Real-time quantitative PCR (RT-PCR) for *MSI2* expression

Total RNA was obtained on day 5 of HSPC culture (72 hrs following CRISPR/Cas9 editing or dCas9-KRAB targeting) using the Total RNA Purification Micro Kit (Norgen, 35300) as per manufacturer’s instructions, including DNase I digestion (Norgen, 25710). 100 to 500 ng of total RNA was used for reverse transcription using the iScript cDNA synthesis (BioRad, # 1708890) or PrimeScript™ RT Master Mix (Takara, RR036A). The cDNA product was used for realtime PCR analysis using iQ SYBR green supermix (BioRad, #1708882). Three technical replicates were performed for each sample, and the mean value was selected for further analysis. The relative expression of each target gene was first normalized to *GAPDH* housekeeping gene expression and then represented as fold changes (2^-ddCt^) relative to the indicated control conditions.

### Intracellular flow cytometry

MSI2 intracellular protein abundance was quantified on day 5 of culture (72 hrs after editing). HSPCs were harvested, washed once in DPBS, and incubated for 30 mins with different fluorescent-labelled antibodies: 1:40 dilution of anti-human CD34 PerCP-Cyanine5.5 (clone 561, BioLegend, #343612), 1:50 dilution of anti-human CD45RA AlexaFluor-488 (clone HI100, BioLegend, #304114) and 1:100 dilution of anti-human CD90 PE (clone 5E10, BioLegend, #328110). Cells were washed, fixed with 4% paraformaldehyde for 15 mins at RT, washed, permeabilized with 0.2% Tween 20 for 15 mins at RT and washed again. Next, cells were then stained with a 1:100 dilution of rabbit anti-MSI2 monoclonal antibody (clone EP1305Y, Abcam, ab76148) for 30 mins at RT, washed, then incubated with a 1:2,000 dilution of goat anti-rabbit AlexaFluor-488 antibody (Invitrogen, A-11034) for 30 mins at RT and washed. Immunophenotypic analyses were performed to quantify the geometric mean fluorescence intensity (MFI) of GFP within CD34^+^CD45RA^-^CD90^+^ gated HSC-enriched cells. Analyses were performed on LSRFortessa (BD Biosciences). Data were analyzed using the FlowJo software.

### Flow cytometry and cell sorting

Immunophenotypic analyses for long-term (LT)-HSCs were performed on adult CD34^+^ HSPCs (on day 8 of serum-free culture, 6 days after editing) or CB CD34^+^ HSPCs (on day 14 of cytokine-free culture, 11 days after editing). Cells were harvested, washed and incubated for 30 mins with fluorescent-labelled antibodies: 1:40 dilution of anti-human CD34 PerCP-Cyanine5.5 (clone 561, BioLegend, #343612), 1:50 dilution of anti-human CD45RA AlexaFluor-488 (clone HI100, BioLegend, #304114), 1:100 dilution of anti-human CD90 PE/Cy7 (clone 5E10, BD Biosciences, # 561558), 1:100 dilution of anti-human CD201 (EPCR) PE (clone RCR-401, BioLegend, #351904), 1:33 dilution of anti-human CD49c (ITGA3) APC (clone ASC-1, BioLegend, #343808). Adult-derived HSPCs in serum-free culture were additionally stained with 1:20 dilution of anti-human CD133 Super Bright 436 (clone 7, BioLegend, #372808). All analyses were performed on LSRFortessa (BD Biosciences). Data were analyzed using the FlowJo software. Total LT-HSC numbers were calculated as a product of the frequency of LT-HSCs by flow cytometry and total cell number in culture.

Fluorescence-activated cell sorting (FACS) was conducted to enrich for HSCs, for DNA editing analysis, RT-PCR of *MSI2* expression, and for single-cell and bulk RNA-sequencing experiments. FACSymphony S6 Cell Sorter (BD Biosciences) was used to sort with a 100 µm nozzle for viable cells (negative for DAPI) with surface expression of CD34^+^CD45RA^-^CD90^+^ markers.

### Xenotransplantation and animal models

All animal procedures were performed under a protocol approved by the Boston Children’s Hospital Institutional Animal Care and Use Committee (IACUC). CD34^+^ HSPCs purified from human cord blood of 2 distinct donors were edited (at AAVS1, ENH-1 or MSI2-KO) on day 2 of culture, and cultured for 48 hr. On day 4 of culture, flow cytometry was used to assess % CD34^+^ in culture for each condition, and 200,000 CD34^+^ cells per mouse were injected via tail vein into Kit-mutant and immunodeficient NOD.Cg-Kit^W41J^Tyr+ Prkdc^sci-d^Il2rg^tm1Wjl^/ThomJ (NBSGW) mice (JAX#026622). To prevent infections, the mice were provided with autoclaved sulfatrim antibiotic water, which was changed weekly. At 16 weeks post-transplantation, the animals were euthanized, and their bone marrows (BMs) and spleens were collected for analysis. BM cells were obtained by flushing the bilateral femurs and tibias, while spleens were carefully minced. Cell suspensions were counted prior to further analyses.

For immunophenotypic analyses of cells retrieved from BM and spleen of xenotransplanted mice, cells were stained with different fluorescent-labelled antibodies: 1:200 dilution of anti-human CD45 APC (clone HI30, BioLegend, #304037), 1:400 dilution of anti-mouse CD45 FITC (clone 30-F11, BioLegend, #103018), 1:400 dilution of anti-human CD3 BV786 (clone SK7, Fisher Scientific, BDB563800), 1:100 dilution of anti-human CD19 PerCP-Cyanine5.5 (clone HIB19, BD Biosciences, #561295), 1:100 dilution of anti-human CD34 APC-Cyanine7 (clone 561, BioLegend, #343614, 1:100 dilution of anti-human CD33 PE (clone HIM3-4, Invitrogen, #12-0339-42), and 1:100 dilution of anti-human CD15 AlexaFluor700 (clone HI98, BioLegend, #301920). Analyses were performed on LSRFortessa (BD Biosciences). Data were analyzed using the FlowJo software.

### Colony-forming unit cell assay

Human CD34^+^ cells were enriched from BM of mice 16-weeks post-transplantation using the Human CD34 MicroBead Kit (Miltenyi Biotec, 130-046-702). 5,000 CD34^+^ HSPCs from each mouse were plated in 1 mL methylcellulose medium (H4434, StemCell Technologies) according to manufacturer’s instructions, with 2 technical replicates per mouse, and grown at 37 °C with 5% CO_2_. Two weeks post-plating, primary colonies were counted, and erythroid, myeloid and mixed colonies were identified according to morphological criteria.

### MSI2-HyperTRIBE

CD34^+^ HSPCs from cord blood were purified using the CD34 MicroBead Kit (Miltenyi Biotec, 130-046-703) according to the manufacturer’s instructions. Purified CD34^+^ cells were cultured for 2 days in IMDM supplemented with 20 % BIT 9500 (StemCell Technologies, #09500), 1X penicillin/streptomycin (Corning, 30-003-CI), and cytokines including 100 ng/mL Human SCF (Peprotech, 300-07-10UG), 10 ng/mL hFLT3L (Peprotech, 300-19-10UG), 100 ng/mL hTPO (Peprotech, 300-18-10UG), and 20 ng/mL hIL-6 (Peprotech, 200-06-20UG). Cells were then transduced with high-titer, concentrated retroviral suspensions encoding either MIG (empty vector control) or MSI2-ADAR constructs in the presence of 8 µg/mL polybrene (Millipore, TR1003G), and spinfected at 700 g for 1.5 hrs. A second round of transduction was performed the following day. GFP-positive cells were sorted using a BD FACSAria cell sorter 24 hrs after second transduction. Three independent biological replicates were performed. RNA was extracted from bulk sorted cells using the SMARTer RNA extraction method. cDNA was subjected to automated paired-end library construction for sequencing on an Illumina HiSeq 2000 platform (PE100 read length). Sequencing was performed at a depth of 30–40 million reads per sample.

### MSI2-ADAR editing site calling

Sequencing reads from three MSI2-ADAR and three control (empty vector) samples were aligned to the human T2T reference genome chm13v2.0^75^ using minimap2 (v.2.26)^76^. All possible single-nucleotide mismatches were piled up with cellsnp-lite (v1.2.3)^77^ in mode 2b in each sample individually, requiring a minimum coverage of 10 and a minor allele frequency of at least 0.01. Mismatches were annotated against the GENCODE v35 GTF to determine their overlap with gene regions. Those annotated as common or rare single-nucleotide polymorphisms (SNPs) in human dbSNP (v155) were excluded from further analysis. Because editing sites in MSI2-ADAR samples may not appear in controls, where they remain unedited and do not appear as mismatches, bcftools (v1.20)^78^ was used to extract allele counts for any mismatch present in at least two MSI2-ADAR samples, across all six samples. We retained only A>G mismatches on the forward strand or T>C mismatches on the reverse strand in control samples to ensure that A or T was the major allele in the control.

Allele counts of retained mismatches were modelled using a negative binomial generalized linear model (GLM) with a log link (via MASS::glm.nb). Specifically, for each mismatch *i*:

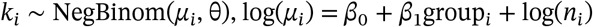

where *k*_*i*_ is the minor allele count, *μ*_*i*_ is the expected count of *k*_*i*_, *θ* is the dispersion parameter estimated by the model, *n*_*i*_ is coverage (with log(*n*_*i*_) serving as an offset), and group_*i*_ indicates whether the sample is MSI2-ADAR or control. P-values of the MSI2-ADAR group coefficient *β*_1_ were adjusted with the Benjamini-Hochberg method. A smaller p-value reflects a larger difference in editing frequency between MSI2-ADAR and control samples. To estimate editing frequency in each group, we regressed the observed edited counts (*k*_*i*_) on the total coverage (*n*_*i*_) without an intercept: *k*_*i*_ ∼ 0 + *n*_*i*_. Ultimately, we kept mismatches that met all of the following criteria: 1) the difference in editing frequencies between MSI2-ADAR and control samples was ≥0.1; 2) the editing frequency in control samples was <0.02; 3) FDR < 0.05 in the GLM test.

The CD34^+^ MSI2-HyperTRIBE score was calculated by taking the sum of total differential editing frequencies per gene (e.g., if a target mRNA was edited at two separate nucleotides in 50% and 70% of reads, then the total score was 1.2). For orthogonal validation, targets and edit sites were presented for MSI2-CLIP-seq in hematopoietic cells reported previously^33^. To identify motifs surrounding MSI2-ADAR editing sites, we used a similar strategy as previously reported^38^. Briefly, sequences extending ±100 bp of each editing site in the 3′ UTR were extracted, and overlapping regions were merged. As a background set, 201-bp segments were randomly selected from 3′ UTRs in the reference genome that did not overlap with the target sequence pool. The HOMER2 software^79^ was used to search for enriched RNA motifs of length 6, 7, or 8.

### Single-cell RNA sequencing and analysis

Droplet-based digital 3’-end single cell RNA sequencing (scRNA-seq) was performed on a Chromium Single-Cell Controller (10X Genomics) using the Chromium Next GEM Single Cell 3’ Reagent Kit v3.1 according to the manufacturer’s instructions. CD34^+^ HSPCs derived from mobilized peripheral blood of 2 distinct healthy donors were edited at AAVS1, with ENH-1 microdeletion or MSI2-KO, on day 2 of culture. 72 hrs after editing, cells were sorted with a 100 µm nozzle for viable cells (DAPI negative) and CD34^+^CD45RA^-^ CD90^+^ cell-surface markers. Approximately 1.6×10^4^ viable cells from each sample were utilized for subsequent scRNA-seq processing, with an estimated recovery of 8,000-12,000 cells per condition. Briefly, single cells were partitioned in Gel Beads in Emulsion (GEMs) and lysed, followed by RNA barcoding, reverse transcription, and PCR amplification (11 cycles). scRNA-Seq libraries were prepared according to the manufacturer’s instructions, checked, and quantified on BioAnalyzer instrument. Sequencing was performed on a Nova Seq S2 (Illumina).

The raw scRNA-seq FASTQ files were processed with the CellRanger (v8.0.1) pipeline to map in the reference genome (GRCh38). We excluded cells with unique molecular identifier (UMI) counts less than 2,000 or greater than 25,000, or mitochondrial UMI fraction higher than 20%, and removed potential doublets by a threshold of doublet score > 0.25 using ScrubletR, which resulted in a total of 57,412 cells for AAVS1 (donor 1 = 10,059 and donor 2 = 12,399), ENH-1 (donor 1 = 7,281 and donor 2 = 8,345), and MSI2-KO (donor 1 = 11,330 and donor 2 = 7,998) edited cells. Symphony R package^60^ was used to project the cells on a human BM reference (https://github.com/andygxzeng/BoneMarrowMap)^80^, to annotate molecularly defined HSCs. A standard Seurat framework (v4.4.0) was used to conduct normalization, principal component analysis (PCA), and dimensionality reduction. The feature-barcode matrix was normalized by the total read count and log-transformed, and the top 3,000 variable features were selected by the vst method in the FindVariableFeatures function. The normalized expression was scaled by Seurat’s ScaleData function, and ribosomal (RPS/RPL) and mitochondrial (MT-) genes were filtered. PCA was performed using the RunPCA function (npc = 30). The sample-dependent technical variation was corrected by using Harmony^61^. Uniform Manifold Approximation and Projection (UMAP) was conducted to reduce dimensions to embed the cells into two-dimensional space. Seurat’s FindMarkers function using “wilcox” method was applied within molecularly defined HSCs to identify differentially expressed genes between ENH-1 vs. AAVS1-edited cells, or MSI2-KO vs. AAVS1-edited cells, with a significance threshold of Benjamini & Hochberg (BH)-adjusted *P* < 0.05, an absolute change in relative expression of ≥ 5%, and minimum percent of expressed cells ≥ 10%. Gene set enrichment analysis was performed using the fGSEA package (https://github.com/ctlab/fgsea/) using the 2024 Hallmark gene sets. Figures were generated using R (v4.4).

### Mapping and validating CHIP-resilience network

The MSI2-DOWN CHIP-resilience network (total 208 genes) was defined by mRNA targets with CD34^+^ MSI2-Hyper-TRIBE score ≥0.2, as well as significantly downregulated following enhancer perturbation (relative expression ≤0.95 and adjusted *P* < 0.05 in ENH-1 vs. AAVS1 HSCs). Ribo-seq data was generated in CD34^+^CD45RA^-^CD90^+^ human HSCs, as previously reported^36^. Raw counts of ribosome protected fragments (RPFs) were normalized to transcript length, and the mean log-normalized RPKM (reads per kilobase per million mapped reads) across three replicates was ranked. GSEA was performed using the fGSEA package. TARGET-seq data (targeted high-sensitivity single cell mutational analysis with parallel RNA-seq) for *TET2* mutant vs. wild-type HSC/MPPs was previously reported^46^. Survival analysis was performed using Survival Genie 2.0^81^, in the TARGET-AML dataset of gene expression in diagnostic BM samples from *N*=1,713 patients^47^. Overall survival from time of diagnosis was stratified by median normalized expression of *MSI2*, or the downstream MSI2-DOWN CHIP-resilience network (*N*=208 genes), with all other default settings.

### Longitudinal clonal growth rate analysis

Targeted, error-corrected sequencing was performed on two serial blood samples from 3,000 people in the Vanderbilt BioVU biobank, with custom-designed probes for 22 CHIP-associated genes, as previously described^82,83^. Vanderbilt University Medical Center’s Institutional Review Board oversees BioVU and approved this project (IRB #201783). UMIs were used for error correction, excluding mutations detected from a single UMI. The mean coverage depth was 1725x after de-duplication. CHIP mutations were called for variants with 100x total read depth, 3 variant allele reads, and variant allele fraction (VAF) ≥ 2% in at least one blood draw. The median time between blood draws was 5.7 years (range: 0.7-13). 657 CHIP mutations (VAF ≥ 2%) were identified at the first time point, of which at the second time point 513 mutations had a VAF ≥ 2% (persistent CHIP) and 144 had a VAF < 2% (transient CHIP). For those with persistent CHIP, growth rate *r* was modelled with a compound interest formula 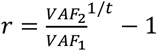 where t is the duration in years between blood draws, as previously done^82,83^. In those with CHIP mutations at baseline, rs17834140 genotypes were extracted using GATK’s HaplotypeCaller^84^. For persistent CHIP mutations, we modeled growth rate *r* ∼ age + age^2^ + VAF + sex + rs17834140-T + driver gene (encoded as a factor), and extracted the coefficient of rs17834140-T. For all CHIP mutations at baseline, we performed logistic regression of transient_CHIP (0/1) ∼ age + age^2^ + VAF + sex + rs17834140-T + driver gene (encoded as a factor). We extracted the odds ratio of transient CHIP by exponentiating the coefficient of rs17834140-T. We repeated the models for the DNMT3A, TET2, ASXL1, and other driver genes separately, and removed driver gene as a covariate for these models.

### Modeling human *ASXL1*-mutant CHIP and bulk RNA-seq

To model germline CHIP-resilience and acquired *ASXL1* mutations, CRISPR/Cas9 was used to edit CD34^+^ cells (from mobilized peripheral blood) on day 1 of culture. HSPCs were edited (as described in previous section) with total 2.0 µL of 100 µM sgRNA, including 1.0 µL to model germline effects [AAVS1: 1.0 µL of sgAAVS1_g1; ENH-1 microdeletion: 0.5 µL each of sgENH-1_g1 and sgENH-1_g2; MSI2-KO: 1 µL of sgMSI2-KO], and 1.0 µL to model somatic mutation [AAVS1: sgAAVS1_g2; ASXL1-exon 12: sgASXL1-ex12]. Immunophenotyping of LT-HSCs was performed at least 7 days after editing.

For bulk RNA-sequencing of ASXL1-edited human HSPCs, FACS was performed on day 8 of culture (7 days after editing) for viable cells (negative for DAPI) and CD34^+^CD45RA^-^CD90^+^ markers to enrich for HSCs. Total RNA was obtained from sorted cells using the Total RNA Purification Micro Kit (Norgen, 35300) as per manufacturer’s instructions. Ultra-low input RNA-seq was conducted with ∼30M reads per sample, across 2 independent experiments in edited HSPCs (AAVS1+AAVS1-sg2, AAVS1+ASXL1, ENH-1+ASXL1 and MSI2-KO+ASXL1). FASTQ files were processed for quality control using FastQC (v0.12.1) to assess sequence quality. Salmon (v1.5.2) was used to align reads to the reference transcriptome (GRCh38, Ensembl v104), to quantify gene expression levels. Transcript abundance estimates were imported into R for differential expression analysis using DESeq2 (v1.34.0). Gene set enrichment analysis was performed using the GO Biological Process 2021 database, and the activated HSC signature previously reported^42^.

### *Asxl1*-mutant mouse model

We crossed Mx1-Cre^-^ *Asxl1*^f/f^ or Mx1-Cre^+^ *Asxl1*^f/f^ mice with Col-tet on-*MSI2*/ROSA-rTTA mice, to generate Mx1-Cre^-^ *Asxl1*^f/f^ *MSI2*^wt/ht^ (Control), Mx1-Cre^-^ *Asxl1*^f/f^ *MSI2*^ht/ht^ (*MSI2*^DOX^), Mx1-Cre^+^ *Asxl1*^f/f^ *MSI2* ^wt/ht^ (*Asxl1*^f/f^) and Mx1-Cre^+^ *Asxl1*^f/f^ *MSI2*^ht/ht^ (*Asxl1*^f/f^+*MSI2*^DOX^) mice. 1 million whole BM cells from donor plus 200K CD45.1 helper BM cells were transplanted into lethally irradiated B6-CD45.1 recipient mice. Depletion of *Asxl1* was initiated one-month post-transplantation by administering three intraperitoneal injections of pIpC HMW (InVivogen, vac-pic) at a dose of 10 mg/kg. MSI2 overexpression was induced by administration of 2 mg/mL Doxycycline hyclate (Fisher Scientific, AC446060050) supplemented with 10 mg/mL sucrose in the drinking water, starting from one month after pIpC treatment. Peripheral blood or BM cells were collected at designated time points and analyzed by flow cytometry.

### Quantification and statistical analysis

In all experiments, data were presented as mean ± standard error of mean. When comparing two samples, a two-tailed Student’s t-test was used to test statistical significance. When one of the two samples was a default value (as in fold change comparison), the one sample t- and Wilcoxon test was applied. When comparing three or more samples, Levene’s test was first used to test equality of variance. If the variance across all samples were tested insignificantly differed, one-way or two-way ANOVA with Dunnett’s test (for multiple comparisons where no reference group is defined) or Tukey’s test (for multiple comparisons where reference group is defined) as post-hoc analysis was used. If the variance across samples was tested to be significantly different, the Kruskal-Wallis test was used instead of ANOVA, with the Dunn test as the post-hoc multiple comparison test. Log-rank test was used for survival analysis. All statistical tests were performed in Graphpad software or R when statistical tests were not available through Graphpad.

**Figure S1:**
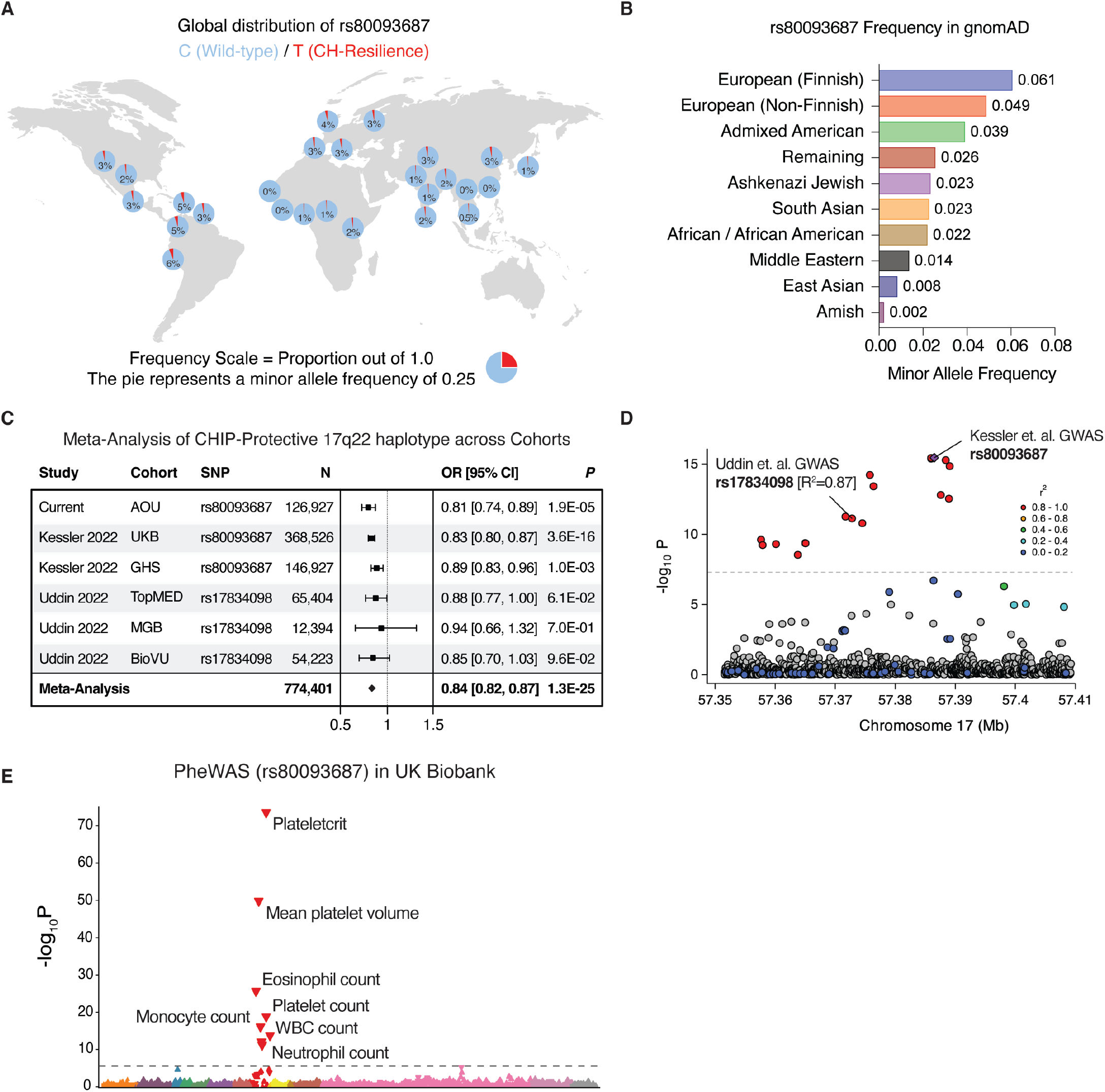
Extended characterization of a CHIP-protective haplotype at the 17q22 locus. **(A)** Global prevalence of rs80093687 across human populations. Figure generated by the Geography of Genetic Variants Browser (*86*). **(B)** Minor allele frequencies of rs80093687 across different ancestries in the gnomAD database. **(C)** Meta-analysis of CHIP odds by rs80093687 or rs17834098 at the 17q22 CHIP-resilience haplotype shows a consistently protective effect across population biobanks. AoU = All of Us; UKB = UK Biobank; GHS = Geisinger Health Study; MGB = Mass General Brigham; BioVU = Vanderbilt University Biobank. **(D)** LocusZoom plot shows high linkage disequilibrium (LD) between rs80093687 and rs17834098 (reported by Uddin *et. al*.^26^ in an independent GWAS for CHIP), suggesting a shared protective haplotype at the 17q22 locus. **(E)** Phenome-wide associations for rs80093687 in UKB^11^.

**Figure S2:**
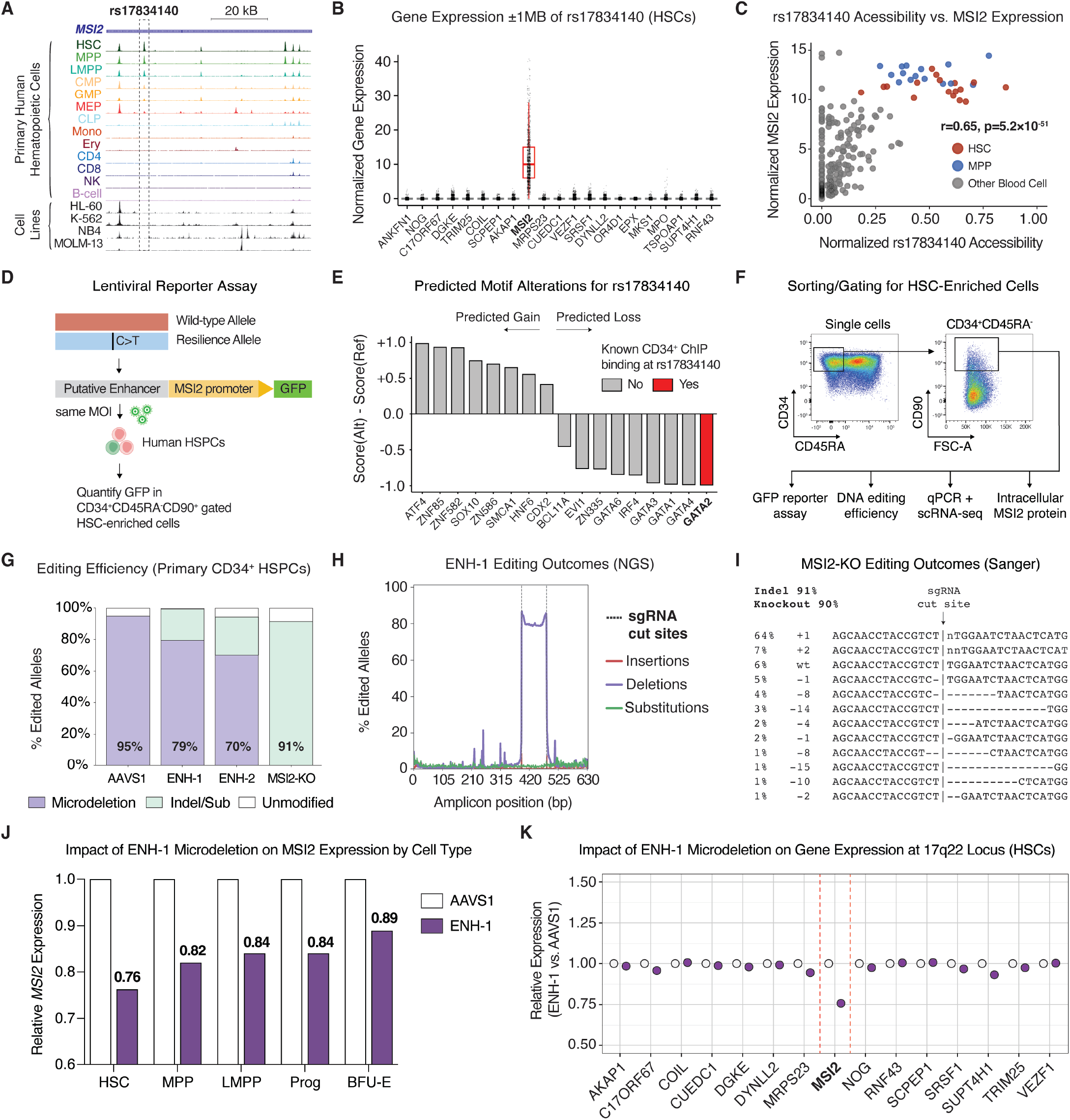
Modeling CHIP-resilience variant effects in primary human HSPCs. **(A)** ATAC-seq tracks at rs17834140 in primary hematopoietic cells and myeloid cell lines, demonstrating selective chromatin accessibility in human hematopoietic stem and early progenitor cells. **(B)** Normalized RNA expression of genes within 1 Mb of rs17834140 in molecularly defined HSCs from human bone marrow. **(C)** Correlation between normalized ATAC-seq reads at rs17834140 and *MSI2* expression across hematopoietic cells (Pearson’s correlation = 0.65). **(D)** Schematic of reporter assay conducted in primary human HSPCs to validate enhancer activity and assess variant effect. **(E)** Predicted gain or loss of putative transcription factor binding sites at rs17834140, using motifBreakR tool^72^, with known binding of factors ascertained from publicly available ChIP-seq datasets in CD34^+^ cells^86^. **(F)** Representative flow cytometry plots showing strategy to either gate or sort CD34^+^CD45RA^-^CD90^+^ HSC-enriched cells for downstream analyses. **(G)** DNA editing efficiencies, inferred through Sanger ICE analysis or next-generation sequencing (NGS), 3 days after editing. **(H)** Editing outcomes at the *MSI2* enhancer by NGS, showing ∼80% efficiency of ∼100bp microdeletion containing rs17834140. **(I)** Editing outcomes at MSI2 exon-targeting knockout (KO) by Sanger sequencing. **(J)** Relative *MSI2* expression 3 days following ENH-1 microdeletion by single-cell RNA-sequencing (scRNA-seq), within molecularly defined HSPC cell types. **(K)** Expression of genes within 1 Mb of rs17834140 in molecularly defined HSCs 3 days after editing at AAVS1 or ENH-1 microdeletion, by scRNA-seq.

**Figure S3:**
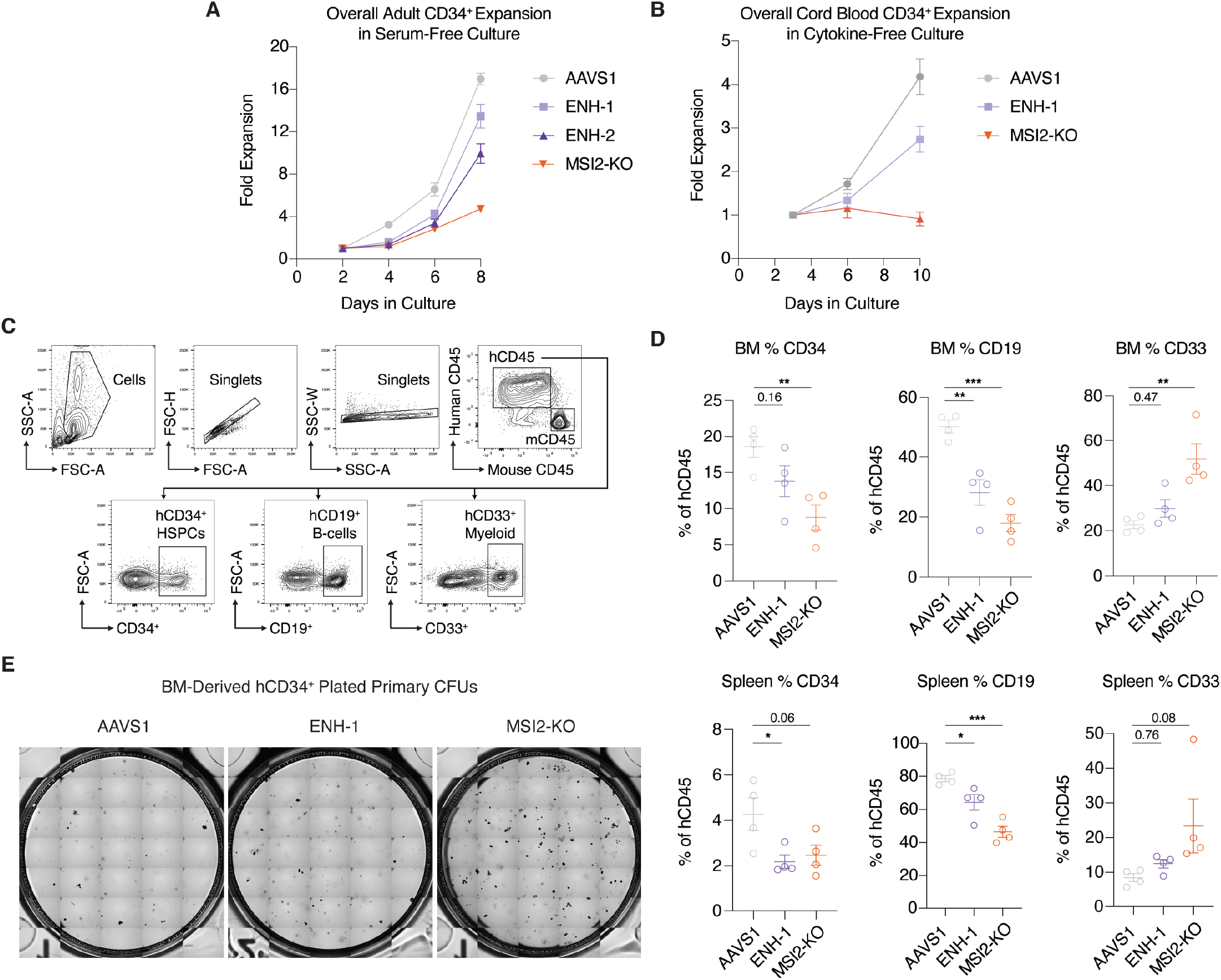
Extended characterization of the *MSI2* enhancer as a regulator of human HSCs. **(A)** Growth curve showing overall fold expansion of adult CD34^+^ human hematopoietic stem and progenitor cells (HSPCs) following editing and culture in serum-free media. **(B)** Growth curve showing overall fold expansion of cord blood CD34^+^ human HSPCs following editing and culture in cytokine-free media. **(C)** Representative flow cytometry gating strategy used to assess human cell engraftment and lineage composition in bone marrow (BM) and spleen of mice 16-weeks after xenotransplantation. **(D)** Proportions of human CD45^+^ cells expressing lineage markers CD34 (HSPCs), CD19 (B-cells) or CD33 (myeloid cells), in BM or spleen of mice 16-weeks after xenotransplantation. **(E)** Representative colony forming assays (CFUs) in bone marrow selected human CD34^+^ HSPCs plated at equal numbers and cultured in methylcellulose media for 14 days, showing increased differentiation capacity of ENH-1 and MSI2-knockout edited cells.

**Figure S4:**
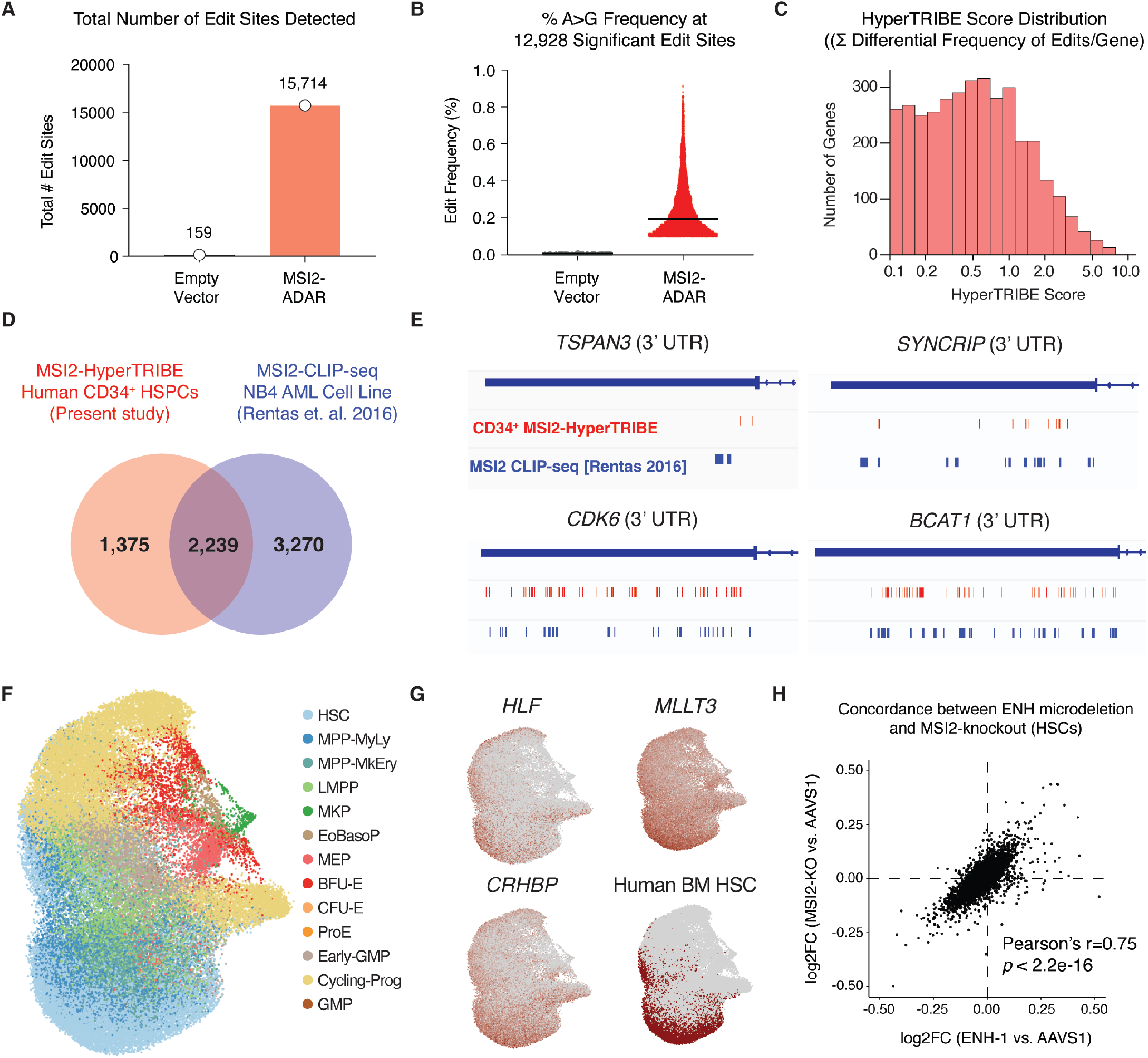
MSI2-HyperTRIBE and single cell profiling of human HSCs. **(A)** Number of A>G edit sites detected in human cord blood CD34^+^ cells transduced with either empty vector or MSI2-ADAR (Adenosine Deaminases Acting on RNA enzyme). **(B)** Editing frequencies at 12,928 significant edit sites in MSI2-HyperTRIBE. **(C)** Histogram showing the distribution of HyperTRIBE scores (sum of all differential editing frequencies per gene). **(D)** Venn diagram showing concordance of CD34^+^ MSI2-HyperTRIBE targets with MSI2-bound mRNAs by CLIP-seq (Cross-Linking Immunoprecipitation Sequencing) in NB4 hematopoietic cell line (reported by Rentas *et. al*.^33^). **(E)** Genome tracks showing concordance of HyperTRIBE editing sites and CLIP-seq reads within 3’ UTR of mRNAs with focal (e.g., *TSPAN3*) or global (e.g., *CDK6*) MSI2 binding. **(F)** Uniform Manifold Approximation and Projection (UMAP) of all CD34^+^CD45RA^-^CD90^+^ HSPCs profiled by single-cell RNA sequencing (scRNA-seq), 72 hrs after editing at AAVS1, ENH-1 or MSI2-knockout. **(G)** UMAPs highlighting molecularly defined human HSCs from a human bone marrow reference map^80^, which are enriched for expression of canonical HSC marker genes. **(H)** Scatter plot showing high correlation between log2FC of differentially expressed genes in ENH-1 vs. AAVS1 and MSI2-KO vs. AAVS1, supporting that MSI2 enhancer microdeletion recapitulate effects of MSI2 loss-of-function edits.

**Figure S5:**
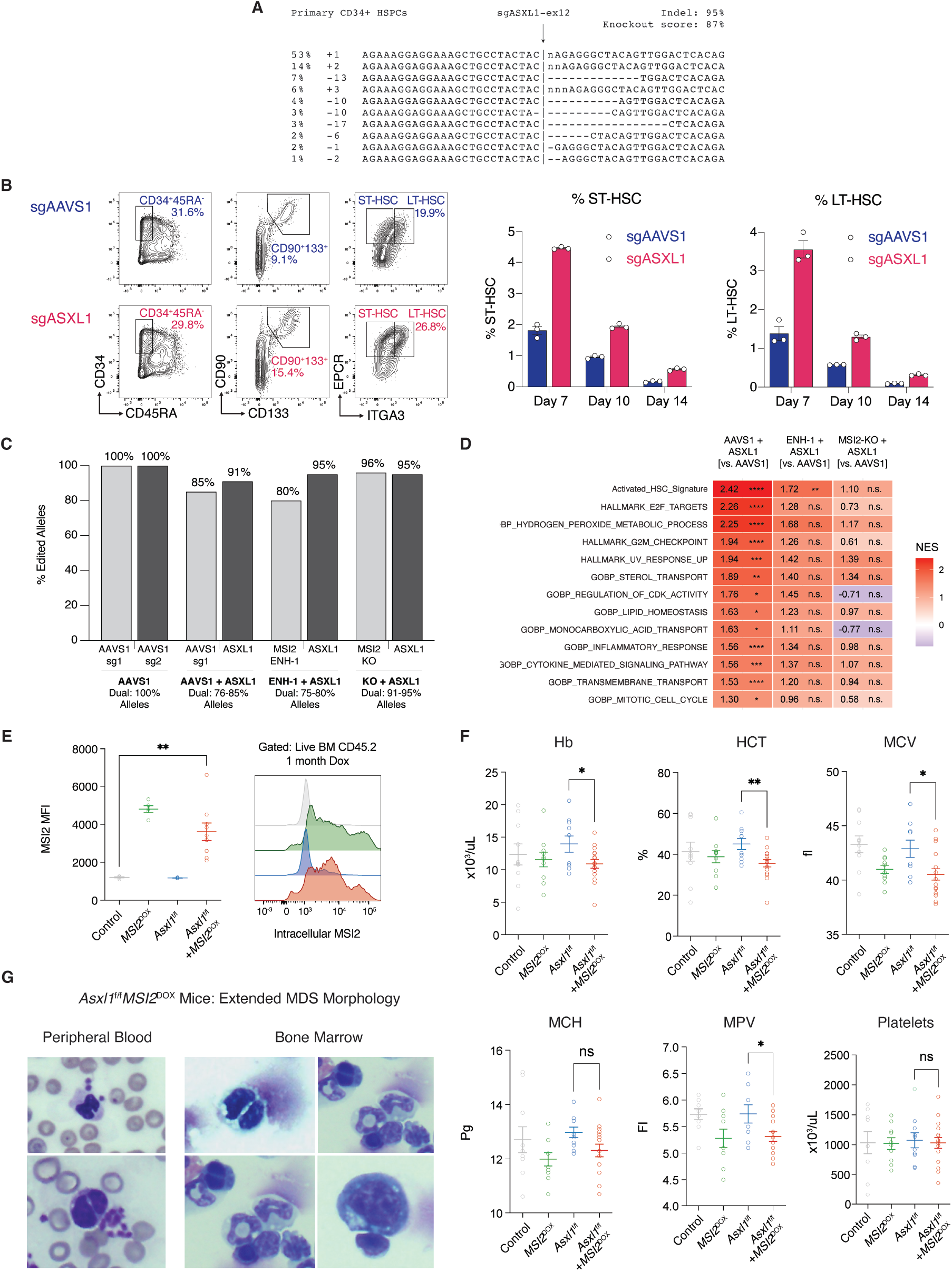
Modeling germline-somatic interactions between MSI2 and ASXL1-CHIP in HSCs. **(A)** Editing outcomes in primary human HSPCs edited with *ASXL1* exon 12 targeting single guide RNA (sgRNA), 7 days post-editing. **(B)** CRISPR model of ASXL1-exon 12 targeting sgRNA expands phenotypic long-term (LT)-HSC maintenance in culture, recapitulating increased stem cell fitness of patient mutations. **(C)** Editing efficiencies at two loci in primary HSPCs dually editing for somatic ASXL1 mutation on a background of the germline resilience effect (AAVS1, ENH-1 or KO), demonstrating that >75% of alleles are dually edited at both loci in culture. **(D)** Gene set enrichment analysis (GSEA) from RNA-seq of CD34^+^CD45RA^-^CD90^+^ HSPCs, showing that MSI2 loss (ENH-1 microdeletion or MSI2-knockout) reverses the transcriptional signature of ASXL1-mutant HSPCs. **(E)** Flow cytometry quantifying MSI2 protein abundance in mouse bone marrow CD45^+^ cells, following *MSI2* induction by 1 month of doxycycline treatment. **(F)** Extended panel of peripheral blood traits in *Asxl1*^f/f^+*MSI2*^DOX^ murine model. Hb = hemoglobin; HCT = hematocrit; MCV = mean corpuscular volume; MCH = mean corpuscular hemoglobin; MPV = mean platelet volume. **(G)** Extended panel of peripheral blood and bone marrow slides demonstrating myelodysplastic syndrome (MDS)-like morphology of *Asxl1*^f/f^+*MSI2*^DOX^ mice.

